# A *Campylobacter* integrative and conjugative element with a CRISPR-Cas9 system targeting competing plasmids: a history of plasmid warfare?

**DOI:** 10.1101/2021.06.01.446523

**Authors:** Arnoud H.M. van Vliet, Oliver Charity, Mark Reuter

## Abstract

Microbial genomes are highly adaptable, with mobile genetic elements (MGEs) such as integrative conjugative elements (ICE) mediating the dissemination of new genetic information throughout bacterial populations. This is countered by defence mechanism such as CRISPR-Cas systems, which limit invading MGEs by sequence-specific targeting. Here we report the distribution of the pVir, pTet and PCC42 plasmids and a new 70-129 kb ICE (CampyICE1) in the foodborne bacterial pathogens *Campylobacter jejuni* and *Campylobacter coli*. CampyICE1 contains a degenerated Type II-C CRISPR system consisting of a sole Cas9 protein, which is distinct from the previously described Cas9 proteins from *C. jejuni* and *C. coli*. CampyICE1 is conserved in structure and gene order, containing blocks of genes predicted to be involved in recombination, regulation, and conjugation. CampyICE1 was detected in 134/5,829 (2.3%) *C. jejuni* genomes and 92/1,347 (6.8%) *C. coli* genomes. Similar ICE were detected in a number of non-jejuni/coli *Campylobacter* species, although these lacked a CRISPR-Cas system. CampyICE1 carries 3 separate short CRISPR spacer arrays containing a combination of 108 unique spacers and 16 spacer variant families. A total of 69 spacers and 10 spacer variant families (63.7%) were predicted to target *Campylobacter* plasmids. The presence of a functional CampyICE1 Cas9 protein and matching anti-plasmid spacers was associated with the absence of the pVir, pTet and pCC42 plasmids (188/214 genomes, 87.9%), implicating that the CampyICE1-encoded CRISPR-Cas has contributed to the exclusion of competing plasmids. In conclusion, the characteristics of the CRISPR-Cas9 system on CampyICE1 suggests a history of plasmid warfare in *Campylobacter*.

**IMPACT STATEMENT:** Understanding pathogen evolution is paramount for enhancing food safety and limiting pathogenic disease in humans and animals. *Campylobacter* species comprise a group of human and animal pathogens with a remarkable success rate, being the most frequent cause of bacterial food-borne disease in high-income countries. A common theme among *Campylobacter* evolution is genomic plasticity, and a significant proportion of this plasticity is driven by horizontal gene transfer (HGT) that results in acquisition of complex traits in one evolutionary event. Understanding the mechanisms of transfer of MGEs and how MGEs such as integrative conjugative elements (ICE) exclude other MGEs is fundamental to understanding *Campylobacter* evolution. CRISPR-Cas9 proteins play a role in bacterial immune systems, mediating the defence against bacteriophage, plasmids, and integrative elements. The use of CRISPR-Cas by a mobile element to fight off competing elements, possibly to the advantage or detriment to their host, also increases our understanding of how important selfish genomic islands undergo co-evolution with bacterial pathogens, and generates insight into the complex warfare between MGEs.

**DATA STATEMENT:** All genome sequences used in this study are available on the National Center for Biotechnology Information (NCBI) Genome database or in the Campylobacter PubMLST website; the assembly accession numbers (NCBI Genome) or genome ID numbers (Campylobacter PubMLST) are listed in Table S1 (available in the online version of this article). Genome assemblies were quality checked based on N_50_, L_50_, genome size and number of contigs. CRISPR Spacer sequences and predicted targets, Cas9 alignments, presence of mobile elements and plasmids are all included in the Supplementary Information.

## INTRODUCTION

The genus *Campylobacter* is a member of the Epsilonproteobacteria, and comprises gram-negative bacteria that are commonly found in the intestines of warm-blooded animals. The best studied members are *C. jejuni* and *C. coli*, which are closely related thermophilic species commonly found in birds and animals involved in agriculture, i.e. poultry, cattle and pigs, while they are also found in many wild birds [1, 2]. They jointly represent the most common bacterial human diarrhoeal pathogens in the developed world, with transmission often foodborne via undercooked meat and cross-contamination in kitchen environments [3, 4]. Other related *Campylobacter* species include the recently described *C. hepaticus* found in poultry [5], *C. upsaliensis* which is a zoonotic *Campylobacter* species from dogs and cats [6], and the *C. lari* group consisting of several species isolated from birds and animals connected to coastal environments [7].

Horizontal gene transfer (HGT) plays a major role in the evolution of microbial genomes [8]. Phages and plasmids are contributors to HGT-driven genomic plasticity, with transfer conducted by either transduction or conjugation, or alternatively by natural transformation [9]. One class of mobile genetic elements (MGE) are the integrative and conjugative elements (ICE), which are self-transferable elements that can mediate excision, form a circular intermediate and often encode the genes for the Type IV conjugative pili used to transfer to a new recipient host cell [10, 11]. ICEs often contain genes required for reversable site-specific recombination, conjugation and regulation, but also carry “cargo” genes that may confer antimicrobial resistance, virulence properties or metabolic capabilities to recipient cells [12], as well as addiction modules ensuring stable maintenance within the host cell [13].

Although acquisition of new genetic traits via HGT may have significant benefits for the recipient cell, the newly acquired sequences can also be detrimental to the host. Therefore cells have developed a diverse set of mechanisms to control entry, integration and expression of foreign DNA [14]. One such system is the Clustered Regularly Interspaced Short Palindromic Repeats (CRISPR) and proteins encoded by CRISPR-associated (Cas) genes, which encode the components of an RNA-guided, sequence-specific immune system against invading nucleic acids, often phages, plasmids and other transferable elements [15]. Many CRISPR-Cas systems have the Cas1 and Cas2 proteins mediating spacer acquisition [16] and other Cas proteins involved in expression, maturation/processing and targeting and interference of the foreign DNA or RNA sequences, commonly phages and plasmids [17]. The RNA-guided endonuclease of the Type II CRISPR-Cas system is the Cas9 (Csn1/Csx12) protein, which mediates processing of CRISPR RNAs and subsequent interference with the targets, in combination with a guide RNA called trans-activating CRISPR RNA (tracrRNA) [18].

Early studies using multilocus sequence typing (MLST) indicated a high level of genetic variability in *Campylobacter* species such as *C. jejuni* and *C. coli* [19], and subsequent comparative genomic analyses have shown that this level of genetic variability is achieved by differences in genetic content and high levels of allelic variability [20–22], likely supported by the natural competence of many *Campylobacter* species. Along with a variety of small plasmids (<10 kb), there are three major classes of 30-60 kb plasmids in *C. jejuni* and *C. coli* (pVir, pTet and pCC42) [23–25], although these are of variable size and gene content [26]. There are also four chromosomally located MGEs first identified in *C. jejuni* RM1221 [27], of which CJIE1 is a Mu-like prophage, CJIE2 and CJIE4 are related temperate prophages [28–31], and CJIE3 is a putative ICE which can contain the *Campylobacter* Type VI secretion system (T6SS) [32, 33].

In a previous study, we showed that 98% of *C. jejuni* genomes investigated contained a Type II-C CRISPR-Cas system consisting of *cas9*-*cas1*-*cas2* genes and a relatively short spacer array (4.9 ± 2.7 spacers, N=1,942 genomes) [34]. In contrast, only 10% of *C. coli* genomes contained a copy of the *C. jejuni* CRISPR-Cas system, while genomes from non-agricultural (environmental) *C. coli* isolates contained a closely related, but separate Type II-C CRISPR-Cas system with the full complement of *cas9*-*cas1*-*cas2* genes, or an orphan *cas9* gene without *cas1* or *cas2* genes [34]. We have expanded this survey of CRISPR-Cas systems in *C. jejuni* and *C. coli*, and show that there is a third, clearly distinct CRISPR-Cas system in both *C. jejuni* and *C. coli*, which is located on a relatively conserved chromosomally located ICE (CampyICE1), and have investigated a possible role of this CRISPR-Cas system in contributing to plasmid competition in *Campylobacter*.

## MATERIALS AND METHODS

### Identification of CRISPR-Cas systems

A collection of complete and draft genome sequences of *C. jejuni* (N=5,829) and *C. coli* (N=1,347) (Table S1) were obtained from the NCBI Genomes database (http://www.ncbi.nlm.nih.gov/genome/browse/) and the *Campylobacter* pubMLST website (http://pubmlst.org/campylobacter/) [35]. Genome assemblies were quality checked based on N_50_, L_50_, genome size and number of contigs, and have been used previously for studying gene distribution in *Campylobacter* [36, 37]. Genome sequences for non-jejuni/coli *Campylobacter* species such as *C. hepaticus*, *C. lari* group and *C. upsaliensis* were obtained from the NCBI genome database using ncbi-genome-download version 0.2.11 (https://github.com/kblin/ncbi-genome-download/). Genome sequences were annotated with Prokka version 1.13 [38], and the annotation searched for Cas9 orthologs using the *C. jejuni* Cj1523c (Cas9) amino acid sequence using BLASTP, while genome sequences were searched using TBLASTN to identify inactivated copies of *cas9* genes. CRISPR arrays were identified as described previously [34], using the CRISPRfinder software (http://crispr.u-psud.fr/Server/) [39] and the CRISPR Recognition Tool CRT [40], further supported by BLAST searches and manual curation. Conservation of sequences was represented using Weblogo [41].

### Prediction of putative targets of CRISPR spacers

A total of 108 unique and 16 variant families of the CampyICE1 CRISPR spacer sequences were used as query on the CRISPRTarget website (http://brownlabtools.otago.ac.nz/CRISPRTarget/crispr_analysis.html) [42], and used to search the Genbank-Phage, Refseq-Plasmid, and Refseq-Viral databases. Only *Campylobacter* targets were included for further analysis. Hits with plasmids from the pVir, pTet and pCC42 families were recorded. Individual genomes with plasmid-specific spacers and positive for either pVir, pTet or pCC42 were searched for the target sequences of that genome using BLAST.

### Analysis of MGE and plasmid distribution

Genome sequences were screened using Abricate (https://github.com/tseemann/abricate) version 0.9.8, with each mobile element/plasmid subdivided into 600 nt fragments used as individual queries, and each 600 nt query sequence was only scored positive with a minimum coverage of 70% and minimum sequence identity of 80%. The CJIE1, CJIE2, CJIE3 and CJIE4 elements were obtained from *C. jejuni* reference strain RM1221 [27]. Nucleotide positions in the RM1221 genome (accession number CP000025) were 207,005-244,247 (CJIE1), 498,503-538,770 (CJIE2), 1,021,082-1,071,873 (CJIE3), and 1,335,703:1,371,932 (CJIE4). The T6SS genes were taken from *C. jejuni* 108 (accession number JX436460). For the CampyICE1 element, genome sequences were screened with the CampyICE1 element from *C. jejuni* strain CCN26 (accession number NZ_FBML01, nucleotide positions contig 11: 109,469-134,196 and reverse strand contig 17: 19,482-78,836), the Clade 1a *C. coli* strain RM1875 (accession number CP007183, nucleotide positions 1,235,330-1,320,414) and the *C. coli* Clade 2 strain C8C3 (accession number FBQX01, nucleotide positions 905,906-996,822). The pCC42 plasmid sequence was obtained from *C. coli* 15-537360 (accession number CP006703), whereas the pTet (accession number CP000549) and pVir (accession number CP000550) plasmid sequences were obtained from *C. jejuni* 81-176. Other plasmids used were pRM3194 (accession number CP014345), pHELV-1 (accession number CP020479) and pSCJK2-1 (accession number CP038863). Genomes were scored as positive for a mobile element or plasmid if >50% positive for 600 nt queries. Samples scoring between 30-50% were manually inspected for distribution of matches and given a final score. Clinker version 0.0.20 [43] was used to generate comparative gene maps of MGE and plasmids, using the default settings. Table S1 includes the presence/absence information of the pCC42, pTet and pVir plasmids, and the CJIE1, CJIE2, CJIE3 and CJIE4 MGE.

### Phylogenetic trees

Core genome MLST allelic profiles were generated for the 5,829 *C. jejuni* and 1,347 *C. coli* genomes using a 678 gene set described previously [44]. Allele calling was performed using chewBBACA version 2.6 [45] using the default settings. The phylogenetic trees were generated using GrapeTree version 1.5.0 [46] with the RapidNJ implementation of Neighbor-Joining, and annotated using the standard 7-gene MLST clonal complexes as determined using the MLST program version 2.19 (https://github.com/tseemann/mlst).

Cas9 protein sequences were aligned with MEGA7 using the MUSCLE algorithm with the default settings [47], and phylogenetic trees constructed using the MEGA7 Neighbor-joining option, pairwise deletion and the Jones-Taylor-Thornton (JTT) model, with 500 bootstraps. Trees were visualised using MEGA7 [47] and Figtree version 1.4.2 (http://tree.bio.ed.ac.uk/software/figtree/).

## RESULTS

### *Campylobacter jejuni* and *C. coli* contain a third type II-C Cas9-encoding gene

A collection of 5,829 *C. jejuni* and 1,347 *C. coli* genomes was searched for the presence of Cas9 orthologs using the *C. jejuni* NCTC11168 Cj1523c and *C. coli* 76639 BN865_15240c amino acid sequences, representative of the two type II-C Cas9 proteins previously detected in *C. jejuni* and *C. coli* [34]. In addition to the *cas9* genes representative of the *C. jejuni*/agricultural *C. coli* and the non-agricultural *C. coli* genomes, a third *cas9* gene was detected in 134 (2.3%) of *C. jejuni* genomes and 92 (6.8%) of *C. coli* genomes, predicted to encode a 965 amino acid protein, with 4 *C. jejuni* and 3 *C. coli* genomes containing an interrupted *cas9* gene. This new *cas9* gene did not have adjacent *cas1* or *cas2* genes. Alignment of the predicted new Cas9 proteins from *C. jejuni* and the *C. coli* clades with Cas9 proteins from members of the genera *Campylobacter* and *Helicobacter* showed that the new Cas9 proteins form a separate cluster (Fig. 1), suggesting these have originated from a more distant common ancestor. Alignment of the additional Cas9 proteins from *C. jejuni* and the different *C. coli* genetic clades showed that the three RuvC motifs, the HNH motif and R-rich region were all conserved (Fig. S1).

**Figure 1.**
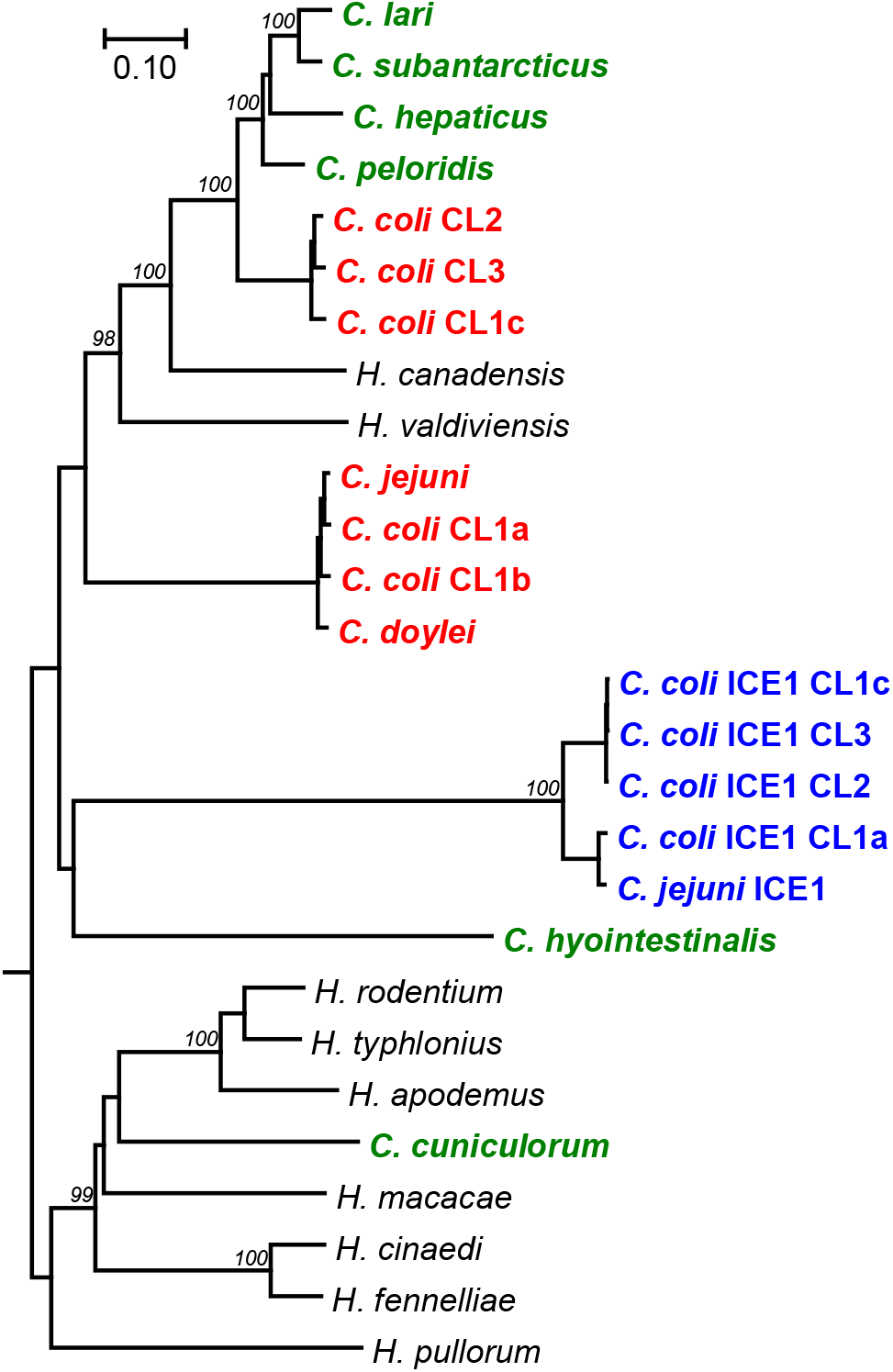
Phylogenetic tree comparing the CampyICE Cas9 proteins with other *Campylobacter* and *Helicobacter* Cas9 proteins. The CampyICE1 Cas9 protein (blue) is distinct from the previously described Cas9 proteins of *C. jejuni* & *C. coli* (red), other *Campylobacter* spp. (green), and selected *Helicobacter* spp. (black). *C. jejuni* subsp. *doylei* is shown as *C. doylei*. The tree was drawn using the Neighbor-Joining method based on an alignment with the MEGA7 Muscle plugin. Bootstrap values are indicated at branches which scored >95%, based on 500 iterations using MEGA7, using the JTT matrix and pairwise deletion. The scale bar represents the number of amino acid substitutions per site. An alignment of a subset of Cas9 proteins with domain annotation is provided in Figure S1.

### The novel CRISPR-Cas system is located on an integrative conjugative mobile element

We first looked for the genomic region containing the gene encoding the new Cas9 protein in completed *C. jejuni* and *C. coli* genomes. Only two complete *C. coli* genomes contained the additional *cas9* gene; an inactivated copy of the *cas9* gene was found on the *C. coli* RM1875 genome, while a complete copy of the gene was present in *C. coli* C8C3. The *cas9* gene was flanked by a short CRISPR-repeat region with five to six repeats, similar to the *Campylobacter* repeat lengths reported previously [34]. Investigation of the surrounding genes showed the downstream presence of a putative Type IV conjugative transfer system, with *traG*, *traN*, *traL* and *traE* genes, as well as a *parM* gene encoding the chromosome segregation protein ParM, while upstream of *cas9*, genes annotated as DNA primase, thymidine kinase, XerC tyrosine recombinase, and an integrase were detected, with the integrase flanked by a tRNA-Met gene as integration site (Fig. 2A), thus matching the common components of an ICE [10]. We have named the *cas9*-containing ICE element CampyICE1.

**Figure 2.**
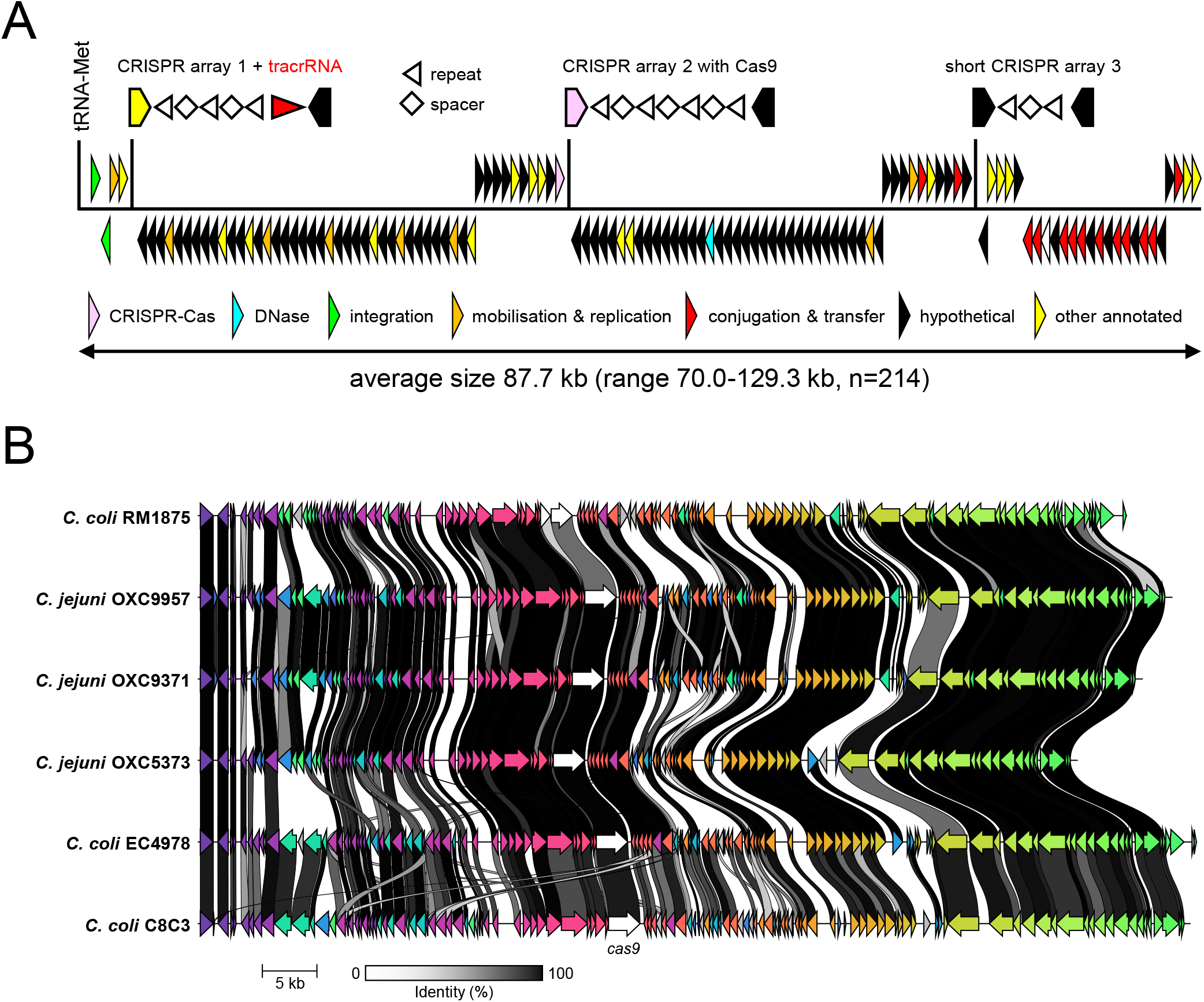
Structure and genetic conservation of CampyICE1 from *C. jejuni* and *C. coli*. **(A)** Schematic overview of the gene structure of CampyICE1 from *C. jejuni* and *C. coli*. The relative positions of the three CRISPR arrays and their transcriptional orientation is shown above the blocks of genes. In the CRISPR arrays, repeats are represented by arrowheads, spacers by diamonds, with the ends of the flanking genes shown. The gene category colors are shown to highlight the large proportion of hypothetical proteins with no known function. **B)** Graphical comparison of CampyICE1 elements from *C. jejuni* and *C. coli* genomes, presented as output of a comparison of Prokka-generated annotations [38] using Clinker [43]. The colours of the arrows in the figure are used to identify homologous blocks of genes, and are not related to the colours used in part A of the figure.

The *C. coli* RM1875 and *C. coli* C8C3 CampyICE1-containing genomic regions were used to search the 134 *C. jejuni* and 92 *C. coli* genomes containing the CampyICE1-*cas9* gene for additional contigs matching the additional CampyICE1 sequences, and ordered these contigs accordingly. We were able to reconstruct the CampyICE1 genomic regions for 81 *C. coli* and 133 *C. jejuni* genomes, annotated these and each showed genetic synteny. The size of the ICE ranged from 70.0-129.3 kb (average 87.7 kb, n=214), and each CampyICE1 region started with a gene encoding a putative integrase (in Genbank often annotated as 30S ribosomal subunit protein), followed by a XerC tyrosine recombinase. There were six relatively conserved blocks of genes downstream, of which the third block ends with the *cas9* gene, and the fourth and the fifth blocks contain genes encoding conjugation proteins (Fig. 2A). Finally, the mobile element also contained up to three putative CRISPR arrays, each with at most a few repeats. The conservation of the CampyICE1 gene synteny is shown in Figure 2B using three *C. coli* and three *C. jejuni* examples.

Searches of the Genbank sequence database for orthologs of CampyICE1 allowed the identification of a similar element in *C. jejuni* subsp. *doylei*, where the element is split into two parts, but lacks the gene block containing the *cas9* gene. There were also regions with sequence and CampyICE1 gene structure similarity in *C. upsaliensis* plasmid pCU110 and *C. iguaniorum* plasmid pCIG1485E, although both lack the *cas9* gene (Fig. S2). Subsequent searches in other *Campylobacter* spp genomes in the Genbank database allowed the identification of other plasmids and potential ICE elements with similar layouts from diverse *Campylobacter* species such as *C. helveticus*, *C. insulaenigrae*, *C. lari* and *C. subantarcticus*, but none of those contained the *cas9* gene (Fig. S2).

### Distribution of CampyICE1 and other mobile elements and linkage to MLST-clonal complexes

To assess whether the distribution of CampyICE1 and other MGEs was linked to specific MLST-types or isolation source, we screened a collection of 5,829 *C. jejuni* and 1,347 *C. coli* genomes [36] using BLAST+ for the presence of CampyICE1, CJIE1, CJIE2, CJIE3, CJIE4, the plasmids pVir, pTet, pCC42, and the CJIE3-associated T6SS (Table 1). The CJIE1 element was the most common in *C. jejuni*, while CJIE4 was the least common of the MGEs from *C. jejuni* RM1221, although still more common than CampyICE1. In *C. coli*, the CJIE1, CJIE2 and CJIE3 elements were present in similar fractions, and again much more common than CJIE4 and CampyICE1 (Table 1). There was clear variation within the CJIE1-CJIE4 genetic elements, mostly in length but also in gene content (Fig. S3), with the CJIE3 element differing due to the presence or absence of the T6SS. With regard to the three plasmids, pVir was rare in both *C. jejuni* and *C. coli*, while pTet is present in approximately a quarter of the *C. jejuni* and *C. coli* genomes. The pCC42 plasmid was relatively rare in *C. jejuni*, but the most common plasmid in *C. coli* (Table 1). The plasmids showed more conservation of gene structure and content (Fig. S4), although there were combinations of plasmids and mobile elements that lead to megaplasmids with phage elements or the T6SS [48] which were not separately included in this analysis.

**Table 1.**
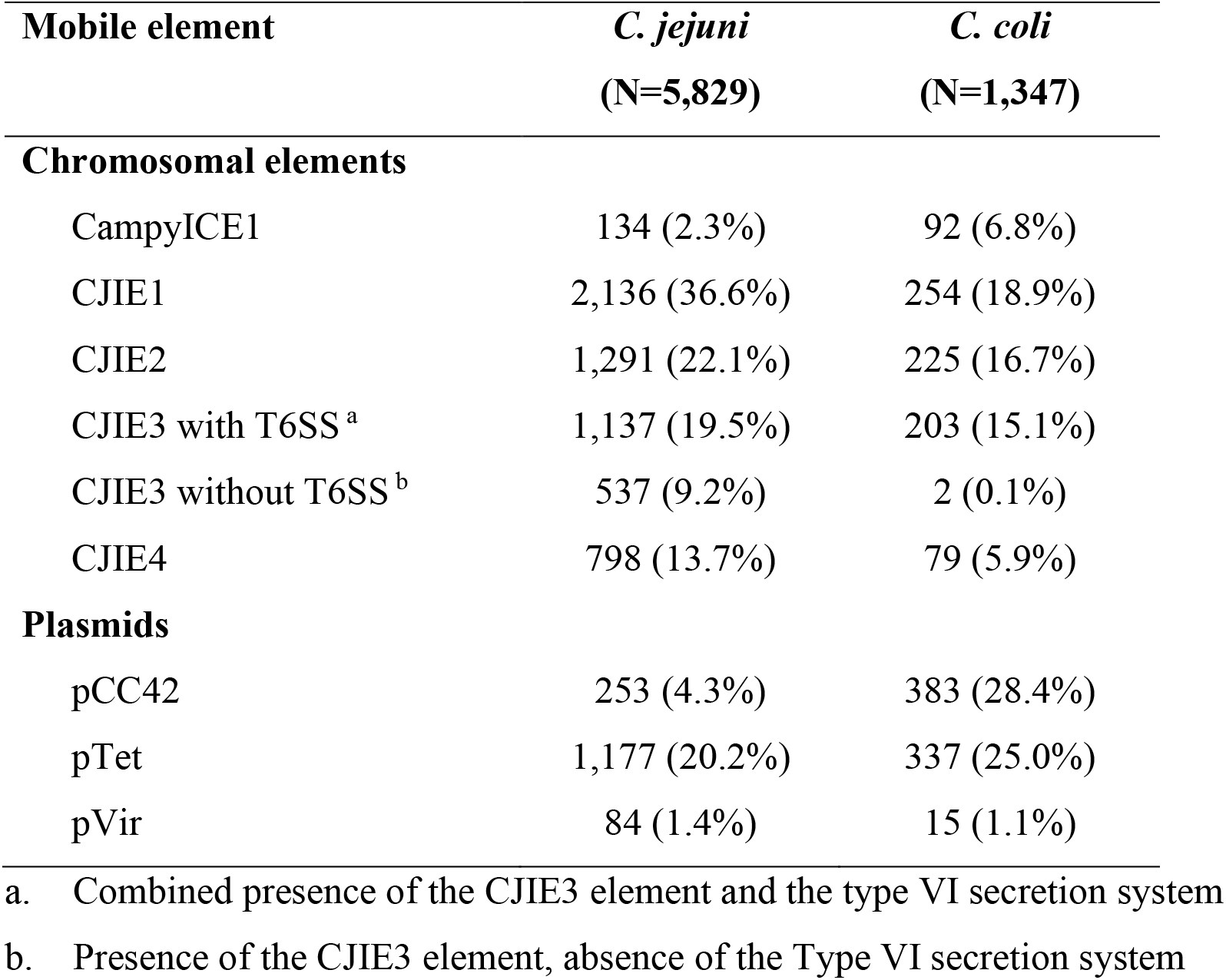
Prevalence of chromosomal and extrachromosomal mobile elements in 5,829 *C. jejuni* and 1,347 *C. coli* genome assemblies.

| Mobile element | *C. jejuni*<br>(N=5,829) | *C. coli*<br>(N=1,347) |
| --- | --- | --- |
| **Chromosomal elements** |  |  |
| CampyICE1 | 134 (2.3%) | 92 (6.8%) |
| CJIE1 | 2,136 (36.6%) | 254 (18.9%) |
| CJIE2 | 1,291 (22.1%) | 225 (16.7%) |
| CJIE3 with T6SS <sup>a</sup> | 1,137 (19.5%) | 203 (15.1%) |
| CJIE3 without T6SS <sup>b</sup> | 537 (9.2%) | 2 (0.1%) |
| CJIE4 | 798 (13.7%) | 79 (5.9%) |
| **Plasmids** |  |  |
| pCC42 | 253 (4.3%) | 383 (28.4%) |
| pTet | 1,177 (20.2%) | 337 (25.0%) |
| pVir | 84 (1.4%) | 15 (1.1%) |

a. Combined presence of the CJIE3 element and the type VI secretion system
b. Presence of the CJIE3 element, absence of the Type VI secretion system

The *C. jejuni* genomes were clustered in a phylogenetic tree based on a 678 gene core genome (cg)MLST scheme [44], which grouped the genomes mostly according to clonal complexes of the seven-gene MLST for *C. jejuni* (Fig. 3) and the different *C. coli* clades (Fig. 4). With the exception of CJIE3 and the associated T6SS in *C. jejuni*, there was no clear association with specific MLST clonal complexes in either *C. jejuni* or *C. coli*. In *C. jejuni*, CJIE3 without the T6SS was restricted to clonal complexes ST-354 and ST-257, while the CJIE3 with T6SS was mostly found in clonal complexes ST-464, ST-353, ST-573 and ST-403 (Fig. 3). There was no obvious link between isolation source and any of the MGEs, although it should be noted that the dataset used is biased towards human isolates. Similar to the mobile elements, the pVir, pTet and pCC42 plasmids did not show an association with either MLST clonal complex in *C. jejuni* or *C. coli* clade, or isolation source (Fig. 3, Fig. 4). The specific distribution per genome is provided in Table S1.

**Figure 3.**
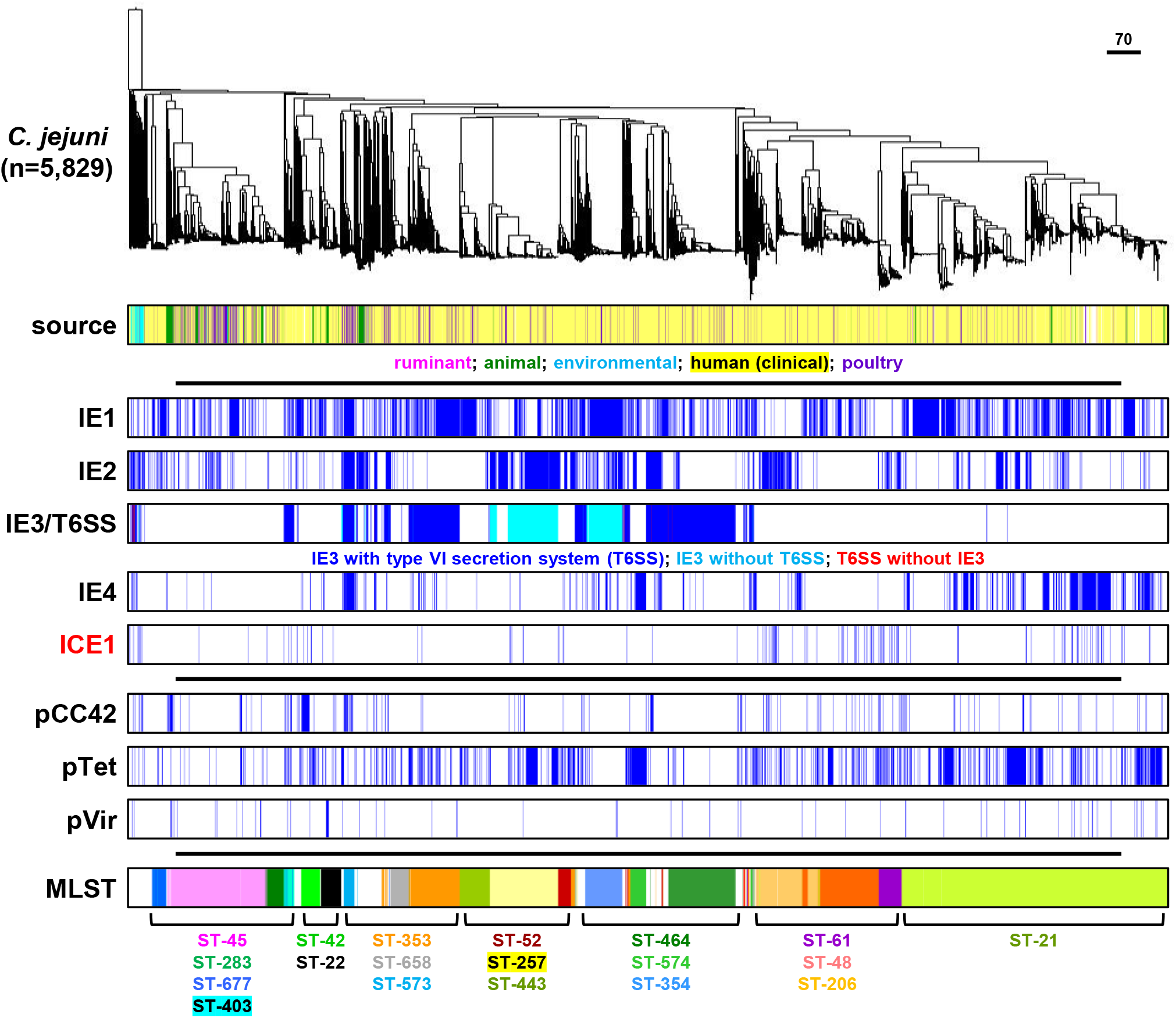
Distribution of mobile elements and plasmids in 5,829 *C. jejuni* genome sequences. The phylogenetic tree was based on core genome MLST. Isolation source category and 7-gene MLST information have been included for comparative purposes.

**Figure 4.**
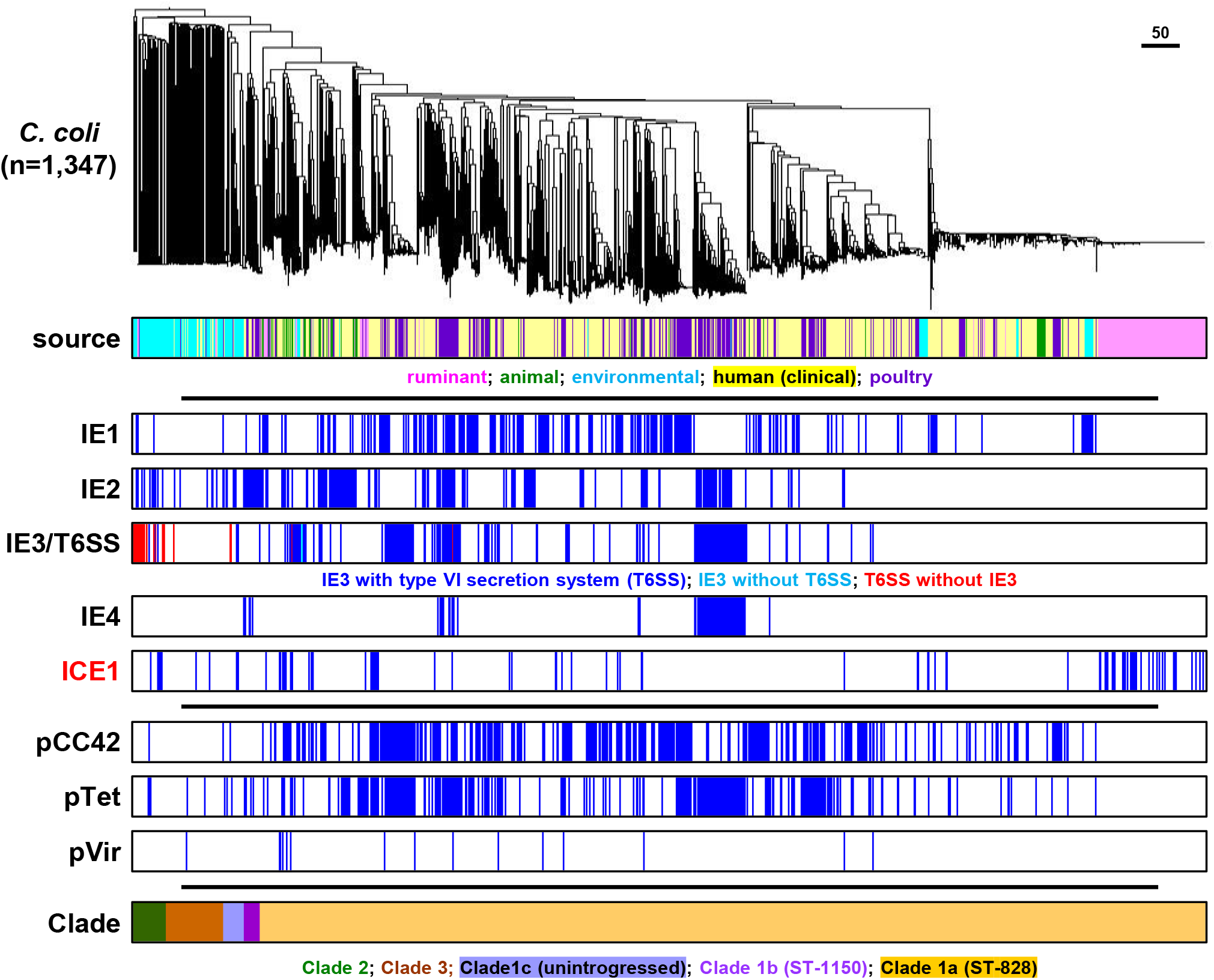
Distribution of mobile elements and plasmids in 1,347 *C. coli* genome sequences. The phylogenetic tree was based on core genome MLST. Isolation source category and 7-gene MLST information have been included for comparative purposes.

### The majority of CampyICE1 CRISPR spacers are predicted to target *Campylobacter* plasmids

CRISPR arrays consist of the CRISPR repeats and the individual spacers, which are used to generate the cRNAs used for interference, and the tracrRNA [18]. The layout of the CampyICE1 CRISPR arrays is distinct from most other Type II CRISPR-Cas systems, where the CRISPR array and tracrRNA are often found directly next to the Cas genes. In contrast, the CampyICE1 system does not contain the ubiquitous *cas1* and *cas2* genes, and has a total of three CRISPR arrays spaced over the element (Fig. 2). We were able to identify spacers from 81 *C. coli* and 133 *C. jejuni* CampyICE1 elements. The first array contained 3.0 ± 1.5 spacers (N=197, range 1-6), and also contained a putative tracrRNA in the opposite transcriptional orientation (Fig. 5A), while the second CRISPR array contained 3.1 ± 1.7 spacers (N=208, range 1-10) and lacked a potential tracrRNA. The third CRISPR array is shorter and contained 1.0 ± 0.6 spacers (N=182, range 1-3). The tracrRNA and repeat sequence are distinct from the previously described *C. jejuni* and *C. coli* CRISPR systems [34], with the changes in the repeat sequence being mirrored in the tracrRNA sequence, thus unlikely to affect functionality (Fig. 5A, 5B). The predicted Protospacer Adjacent Motif (PAM) was 5′-A(C/T)A(C/T) (Fig. 5A), which matches well with the 5′-ACAc PAM-motif described for the *C. jejuni* Cas9 protein [34, 49].

**Figure 5.**
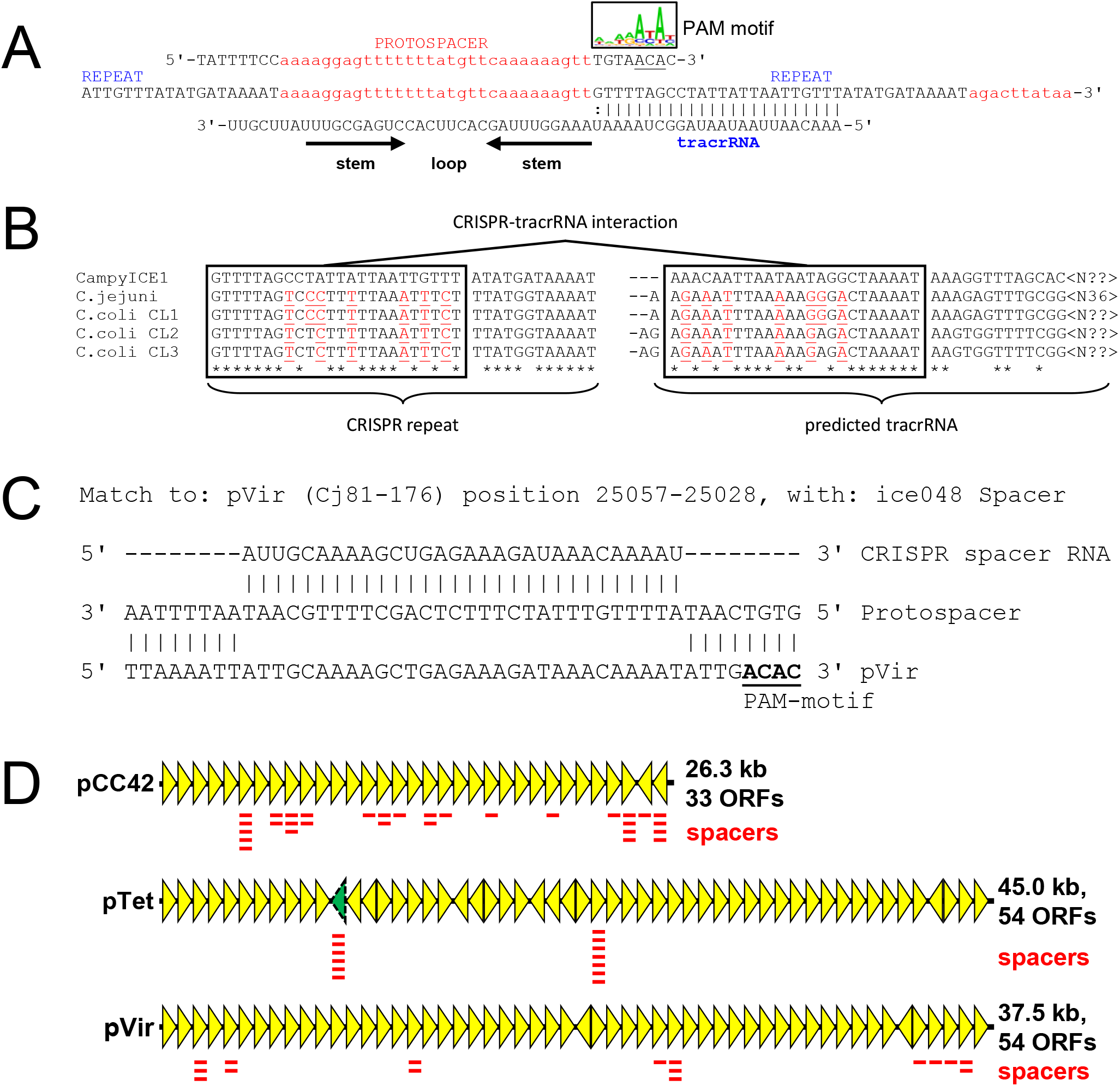
Characteristics of the CampyICE1 CRISPR spacers, protospacers and tracrRNA, and predicted plasmid targeting by the CampyICE1 CRISPR-Cas9 system. (A) A section of the CRISPR array is shown (center) with the corresponding protospacer (top) with 8 nt flanking sequences which contain the PAM motif at the 3′ end of the protospacer, represented using a sequence logo. The tracrRNA sequence and structure are included below. (B) Comparison of the CRISPR-repeats and predicted tracrRNA part of CampyICE1, *C. jejuni* and the three *C. coli* clades. The tracrRNA and CRISPR-repeat show matching changes as indicated by red underlined residues. Asterisks indicate conserved nucleotides, boxes indicate the complementary sequences in CRISPR repeat and tracrRNA. (C) Example of a CampyICE1 CRISPR spacer perfectly matching a segment of the *C. jejuni* 81-176 pVir plasmid. (D) Schematic representation of the pCC42, pTet and pVir family of plasmids (based on the *C. coli* 15-537360 pCC42 plasmid and the *C. jejuni* 81-176 pTet and pVir plasmids), with the locations of plasmid-targeting CampyICE1 spacers indicated. For pTet, the approximate location of target gene YSU_08860 (absent from the *C. jejuni* 81-176 plasmid) is indicated by the dashed, green colored arrowhead. More information on specific spacers and their targets is provided in Table S2.

Comparison of the spacers from 214 CampyICE1 elements showed that these consisted of 108 unique spacer sequences, and an additional 40 spacers that were subdivided in 16 variant families, where 2-6 spacers had one or two nucleotide differences to each other and were predicted to match the same targets (Table S2). The spacers were used to search phage and plasmid databases for putative targets, and a total of 62 unique spacers and eight variant families were predicted to target the *Campylobacter* plasmids pCC42 (31 unique spacers, two variants), pTet (16 unique spacers, six variants) and pVir (15 unique spacers, see Fig. 5C for an example). Furthermore there were spacers predicted to target the *Campylobacter helveticus* plasmid pHELV-1 (one unique spacer) and pSCJK2-1 from *C. jejuni* SCJK2 (six unique spacers, two variants). The pHELV-1 and pSCJK2-1 plasmids were not detected in the 5,829 *C. jejuni* and 1,347 *C. coli* genomes used in this study. The predicted targets on the plasmids pCC42, pTet and pVir were plotted against the plasmid maps (Fig. 5D), and showed that targets for pCC42 and pVir were found in multiple genes on these two plasmids, whereas the targets on pTet were limited to two genes, of which YSU_08860 is not universally present on plasmids of the pTet family (Fig. 5D).

### Plasmid-mapping CampyICE1 CRISPR spacers are associated with an absence of the corresponding plasmids

To assess whether the CampyICE1 CRISPR-Cas9 system can function to exclude plasmid by using plasmid-mapping spacers, the 226 *C. jejuni* and *C. coli* CampyICE1-positive genome assemblies were searched for the presence of plasmid contigs and matches with spacer sequences (Table S3 and Table S4). As one possible escape for CRISPR-Cas9 surveillance could be sequence mutations/changes in the plasmids, we also checked whether the predicted plasmid-matching spacer would recognise any sequence in the genome assemblies (which include plasmid contigs). Of the *C. coli* assemblies, spacers were detected in 81/92 assemblies, and 56 had no plasmid/spacer matches. Of the 25 assemblies where there were plasmid/spacer matches, three had an inactivated CampyICE1 *cas9* gene, and 11 did not have sequences matching the spacer(s) or only partial matches in their genome assembly, suggesting that mutations in the plasmid sequence have made the spacer unusable. This left 11 *C. coli* assemblies with a functional *cas9* gene and spacer matching the pCC42 plasmid. Similarly, for *C. jejuni*, spacers were detected in 133/134 genomes, and 109 had no plasmid/spacer matches. Of the 24 assemblies where there were plasmid/spacer matches, two had an inactivated CampyICE1 *cas9* gene with frameshifts and stop codons, and seven did not have sequences matching the spacer(s) or only partial matches in their genome assembly. This left 15 *C. jejuni* assemblies with a functional *cas9* gene and spacer matching the pCC42 (seven) and pTet (eight) plasmids. The matching of spacers, CampyICE1 Cas9 status and plasmid presence/absence is given in Figure 6, with more detailed data in Table S3 and Table S4.

**Figure 6.**
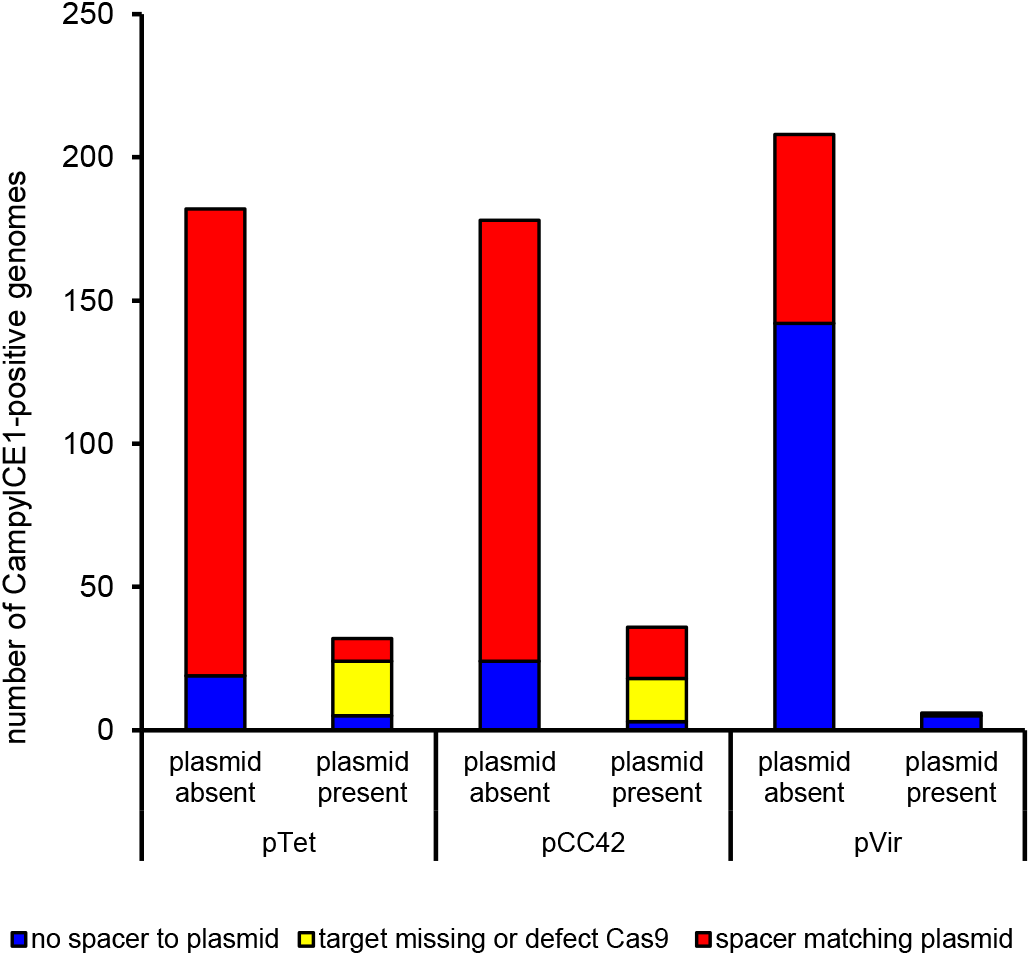
Low prevalence of pVir, pTet and pCC42 plasmids in CampyICE1-positive *C. jejuni* and *C. coli* is associated with CRISPR-spacers targeting these plasmids. The *C. jejuni* and *C. coli* isolates have been combined in this graph; specific data per isolate and spacer are available in Table S3, data for *C. jejuni* and *C. coli* separately are provided in Table S4.

## DISCUSSION

In the last 25 years, CRISPR-Cas has gone from a relatively obscure repeat system in bacteria to a Nobel Prize winning phenomenon [50]. CRISPR-Cas systems are widespread in prokaryotic organisms, and while early reports predicted them to be a bacterial version of the adaptive immune system against phages, it is now clear that they target a wide variety of MGEs, and can also have a diverse set of alternative functions. Recent studies show that CRISPR-Cas systems are not just located on genomes, but can also be found on MGEs. Type IV and Type I CRISPR-Cas systems have been reported on enterobacterial plasmids [51, 52], and have been predicted to function in competition between plasmids [53]. *Vibrionaceae* species contain a variety of CRISPR-Cas systems associated with putative MGEs and genomic islands [54, 55], although data on their potential role in MGE competition are still lacking. To our knowledge, our study is the first to feature an incomplete Type II-C CRISPR-Cas9 system that is associated with an MGE, and where the majority of spacers matched competing plasmids. We have shown that CampyICE1 is highly conserved in both *C. jejuni* and *C. coli*, that it has up to three short spacer arrays on the ICE, and that the presence of a functional CampyICE1 CRISPR-Cas system and anti-plasmid spacers is associated with the absence of the three targeted plasmid types in *C. jejuni* and *C. coli*.

The Type II-C Cas9 protein encoded on CampyICE1 is closely related to the Cas9 proteins found in other *Campylobacter* and *Helicobacter* species, but clusters separately, suggesting it may have been co-opted from a genomic location in an ancestral *Campylobacteraceae* species. Interestingly, CampyICE1 lacks the *cas1* and *cas2* genes [56], a feature which has also been noted for the hypercompact Cas12j (CasΦ) system found on certain bacteriophages [57]. The Cas12j system much resembles the CampyICE1 Cas9 system described here, as they share a limited CRISPR spacer repertoire [58]. The lack of Cas1 and Cas2 components could mean that the CampyICE1 system is incapable of acquiring new spacers, which is supported by the relative lack of spacer diversity in the 214 genomes containing CampyICE1. However, we cannot exclude that the CampyICE1 Cas9 may be able to co-opt the Cas1 and Cas2 proteins from the chromosomal version of the CRISPR-Cas system in *C. jejuni* and *C. coli*, although this is speculative. We have previously shown that ~98% of all *C. jejuni* genomes have a CRISPR-Cas system, while this is more limited in *C. coli*, where only ~10% of *C. coli* genomes have a CRISPR-Cas system [34]. Since the diversity in CRISPR spacers is also low in the chromosomal version of CRISPR-Cas of *C. jejuni* and *C. coli* and most spacers cannot (yet) be linked to mobile elements or phages [34, 59–61], it may represent additional or alternative functions for Cas9 in *C. jejuni*, such as control or activity in virulence [62–66]. However, this is not the case for the CampyICE1 CRISPR-Cas9 system, as a majority of spacers can be linked to the three main families of plasmids in *C. jejuni* and *C. coli*: pTet, pVir and pCC42.

In our collection of genomes, 41.7% of *C. coli* and 24.3% of *C. jejuni* genomes are predicted to contain one or more of these three plasmids, in different combinations. The three plasmids do not show signs of incompatibility, as 93 *C. jejuni* and 166 *C. coli* genomes had a combination of two plasmids or all three plasmids together. The role of these plasmids in *C. jejuni* and *C. coli* is still unclear, but they can carry virulence factors and contribute to the dissemination of antibiotic resistance. However, plasmids are not absolutely required for this, and plasmid-free isolates are also common. This is similar for the CJIE elements, where different combinations of the CJIE-elements and CampyICE1 were detected. The different roles of the CJIE-elements in *C. jejuni* and *C. coli* is still not clear, although the T6SS from CJIE3 has been linked with virulence [32, 33, 67, 68], and the DNases of the CJIE1, CJIE2, and CJIE4 elements are associated with reduced biofilm formation and reduced natural transformation [29, 30, 69].

The CRISPR-Cas9 system of the CampyICE1 element has some unique properties, as there are up to three short CRISPR arrays on the mobile element, with the essential tracrRNA not located with the *cas9* gene but located in another CRISPR spacer array on CampyICE1. Although the arrays detected were small, there were still 108 unique spacers and 16 spacer families, with a spacer family defined as spacers differing by one or two nucleotides only. The majority of CampyICE CRISPR spacers and variants were predicted to target *Campylobacter* plasmids (69 spacers and 10 variants, 63.7%), with most spacers predicted to target pCC42, pTet and pVir, the three major plasmids in *C. jejuni* and *C. coli*, which is a very high proportion compared to many other CRISPR-Cas studies. For example, a study on type IV CRISPR-Cas systems could only match 12% of spacers with targets, and this was reduced to only 7% for the non-type IV CRISPR-Cas systems [53]. In our previous study [34] we were also unable to match most *Campylobacter* spacers with putative targets, which is common. The presence of CampyICE1, functional CRISPR-Cas9 and anti-plasmid spacers was associated with the absence of the competing plasmids targeted, suggesting that CampyICE1 has used its CRISPR-Cas9 system for “plasmid warfare” as a form of incompatibility. The match is not perfect, as there are several examples of a complete CampyICE1 CRISPR-Cas9 system with plasmid-targeting spacers, to which the spacers mapped were present with 100% sequence identity between spacer and predicted plasmid contigs (Table S3, Table S4, Fig. 6). This could potentially mean that the CRISPR-Cas system can prevent acquisition of new plasmids, but for unknown reasons is unable to remove plasmids already present, although this is highly speculative. It also suggests that the CampyICE1 plasmid restriction can be avoided by mutation of the target site disrupting the sequence matching, making the system less functional, especially in a bacterium known for its high levels of genetic variation. We also speculate that DNA modification and transcriptional variation/regulation may play a role in spacer-target discrepancies.

In summary, we have identified a new putative mobile element in *C. jejuni* and *C. coli* that contains a degenerated CRISPR-Cas9 system predicted to employ this CRISPR-Cas system to compete with other families of *Campylobacter* plasmids. We also show that mobile elements and plasmids are semi-randomly distributed within a large set of *C. jejuni* and *C. coli* genomes, and display significant levels of genetic variation within the elements. This fits well with the previously described genetic variability of the genus *Campylobacter*, and adds to the complexity of mobile elements present within these successful foodborne human pathogens.

## ACKNOWLEDGMENTS

This publication made use of the PubMLST website (http://pubmlst.org/) developed by Keith Jolley and sited at the University of Oxford. The development of that website was funded by the Wellcome Trust.

## AUTHOR CONTRIBUTIONS

A.H.M.v.V. conceived the study and study design, performed analysis and wrote the paper; O.C. and M.R. contributed to study design, performed analysis and writing of the paper.

## REFERENCES

1. Rossler E, Signorini ML, Romero-Scharpen A, Soto LP, Berisvil A et al. Meta-analysis of the prevalence of thermotolerant Campylobacter in food-producing animals worldwide. Zoonoses Public Health 2019;66(4):359–369 doi: 10.1111/zph.12558

2. Griekspoor P, Colles FM, McCarthy ND, Hansbro PM, Ashhurst-Smith C et al. Marked host specificity and lack of phylogeographic population structure of Campylobacter jejuni in wild birds. Mol Ecol 2013;22(5):1463–1472 doi: 10.1111/mec.12144

3. Nichols GL, Richardson JF, Sheppard SK, Lane C, Sarran C. Campylobacter epidemiology: a descriptive study reviewing 1 million cases in England and Wales between 1989 and 2011. BMJ Open 2012;2(4):e001179 doi: 10.1136/bmjopen-2012-001179

4. Romdhane RB, Merle R. The Data Behind Risk Analysis of Campylobacter Jejuni and Campylobacter Coli Infections. Curr Top Microbiol Immunol 2021;431:25–58 doi: 10.1007/978-3-030-65481-8_2

5. Crawshaw T. A review of the novel thermophilic Campylobacter, Campylobacter hepaticus, a pathogen of poultry. Transbound Emerg Dis 2019;66(4):1481–1492 doi: 10.1111/tbed.13229

6. Campagnolo ER, Philipp LM, Long JM, Hanshaw NL. Pet-associated Campylobacteriosis: A persisting public health concern. Zoonoses Public Health 2018;65(3):304–311 doi: 10.1111/zph.12389

7. Miller WG, Yee E, Chapman MH, Smith TP, Bono JL et al. Comparative genomics of the Campylobacter lari group. Genome Biol Evol 2014;6(12):3252–3266 doi: 10.1093/gbe/evu249

8. Brockhurst MA, Harrison E, Hall JPJ, Richards T, McNally A et al. The Ecology and Evolution of Pangenomes. Curr Biol 2019;29(20):R1094–R1103 doi: 10.1016/j.cub.2019.08.012

9. Penades JR, Chen J, Quiles-Puchalt N, Carpena N, Novick RP. Bacteriophage-mediated spread of bacterial virulence genes. Curr Opin Microbiol, Review 2014;23C:171–178 doi: 10.1016/j.mib.2014.11.019

10. Johnson CM, Grossman AD. Integrative and Conjugative Elements (ICEs): What They Do and How They Work. Annu Rev Genet 2015;49:577–601 doi: 10.1146/annurev-genet-112414-055018

11. Delavat F, Miyazaki R, Carraro N, Pradervand N, van der Meer JR. The hidden life of integrative and conjugative elements. FEMS Microbiol Rev 2017;41(4):512–537 doi: 10.1093/femsre/fux008

12. Partridge SR, Kwong SM, Firth N, Jensen SO. Mobile Genetic Elements Associated with Antimicrobial Resistance. Clin Microbiol Rev 2018;31(4):e00088–00017 doi: 10.1128/CMR.00088-17

13. Burrus V. Mechanisms of stabilization of integrative and conjugative elements. Curr Opin Microbiol 2017;38:44–50 doi: 10.1016/j.mib.2017.03.014

14. Labrie SJ, Samson JE, Moineau S. Bacteriophage resistance mechanisms. Nat Rev Microbiol 2010;8(5):317–327 doi: 10.1038/nrmicro2315

15. Makarova KS, Wolf YI, Alkhnbashi OS, Costa F, Shah SA et al. An updated evolutionary classification of CRISPR-Cas systems. Nat Rev Microbiol 2015;13(11):722–736 doi: 10.1038/nrmicro3569

16. McGinn J, Marraffini LA. Molecular mechanisms of CRISPR-Cas spacer acquisition. Nat Rev Microbiol 2019;17(1):7–12 doi: 10.1038/s41579-018-0071-7

17. Hille F, Richter H, Wong SP, Bratovic M, Ressel S et al. The Biology of CRISPR-Cas: Backward and Forward. Cell 2018;172(6):1239–1259 doi: 10.1016/j.cell.2017.11.032

18. Makarova KS, Wolf YI, Iranzo J, Shmakov SA, Alkhnbashi OS et al. Evolutionary classification of CRISPR-Cas systems: a burst of class 2 and derived variants. Nat Rev Microbiol 2020;18(2):67–83 doi: 10.1038/s41579-019-0299-x

19. Sheppard SK, McCarthy ND, Falush D, Maiden MC. Convergence of Campylobacter species: implications for bacterial evolution. Science 2008;320(5873):237–239 doi: 10.1126/science.1155532

20. Baig A, McNally A, Dunn S, Paszkiewicz KH, Corander J et al. Genetic import and phenotype specific alleles associated with hyper-invasion in Campylobacter jejuni. BMC Genomics 2015;16:852 doi: 10.1186/s12864-015-2087-y

21. Sheppard SK, Jolley KA, Maiden MC. A Gene-By-Gene Approach to Bacterial Population Genomics: Whole Genome MLST of Campylobacter. Genes (Basel) 2012;3(2):261–277 doi: 10.3390/genes3020261

22. Yahara K, Meric G, Taylor AJ, de Vries SP, Murray S et al. Genome-wide association of functional traits linked with Campylobacter jejuni survival from farm to fork. Environ Microbiol 2017;19(1):361–380 doi: 10.1111/1462-2920.13628

23. Batchelor RA, Pearson BM, Friis LM, Guerry P, Wells JM. Nucleotide sequences and comparison of two large conjugative plasmids from different Campylobacter species. Microbiology 2004;150(Pt 10):3507–3517 doi: 10.1099/mic.0.27112-0

24. Pearson BM, Rokney A, Crossman LC, Miller WG, Wain J et al. Complete genome sequence of the clinical isolate Campylobacter coli 15-537360. Genome Announc 2013;1(6):e01056–01013 doi: 10.1128/genomeA.01056-13

25. Bacon DJ, Alm RA, Burr DH, Hu L, Kopecko DJ et al. Involvement of a plasmid in virulence of Campylobacter jejuni 81-176. Infect Immun 2000;68(8):4384–4390 doi: 10.1128/iai.68.8.4384-4390.2000

26. Marasini D, Fakhr MK. Complete Genome Sequences of Plasmid-Bearing Multidrug-Resistant Campylobacter jejuni and Campylobacter coli Strains with Type VI Secretion Systems, Isolated from Retail Turkey and Pork. Genome Announc 2017;5(47):e01360–01317 doi: 10.1128/genomeA.01360-17

27. Fouts DE, Mongodin EF, Mandrell RE, Miller WG, Rasko DA et al. Major structural differences and novel potential virulence mechanisms from the genomes of multiple *campylobacter* species. PLoS biology 2005;3(1):e15 doi: 10.1371/journal.pbio.0030015

28. Parker CT, Quinones B, Miller WG, Horn ST, Mandrell RE. Comparative genomic analysis of Campylobacter jejuni strains reveals diversity due to genomic elements similar to those present in C. jejuni strain RM1221. J Clin Microbiol 2006;44(11):4125–4135 doi: 10.1128/JCM.01231-06

29. Gaasbeek EJ, Wagenaar JA, Guilhabert MR, van Putten JP, Parker CT et al. Nucleases encoded by the integrated elements CJIE2 and CJIE4 inhibit natural transformation of Campylobacter jejuni. J Bacteriol, Research Support, Non-U.S. Gov’t 2010;192(4):936–941 doi: 10.1128/JB.00867-09

30. Gaasbeek EJ, Wagenaar JA, Guilhabert MR, Wosten MM, van Putten JP et al. A DNase encoded by integrated element CJIE1 inhibits natural transformation of Campylobacter jejuni. J Bacteriol 2009;191(7):2296–2306 doi: 10.1128/JB.01430-08

31. Clark CG, Chen CY, Berry C, Walker M, McCorrister SJ et al. Comparison of genomes and proteomes of four whole genome-sequenced Campylobacter jejuni from different phylogenetic backgrounds. PLoS One 2018;13(1):e0190836 doi: 10.1371/journal.pone.0190836

32. Bleumink-Pluym NM, van Alphen LB, Bouwman LI, Wosten MM, van Putten JP. Identification of a Functional Type VI Secretion System in Campylobacter jejuni Conferring Capsule Polysaccharide Sensitive Cytotoxicity. PLoS Pathog 2013;9(5):e1003393 doi: 10.1371/journal.ppat.1003393

33. Lertpiriyapong K, Gamazon ER, Feng Y, Park DS, Pang J et al. Campylobacter jejuni type VI secretion system: roles in adaptation to deoxycholic acid, host cell adherence, invasion, and in vivo colonization. PLoS One 2012;7(8):e42842 doi: 10.1371/journal.pone.0042842

34. Pearson BM, Louwen R, van Baarlen P, van Vliet AHM. Differential Distribution of Type II CRISPR-Cas Systems in Agricultural and Nonagricultural Campylobacter coli and Campylobacter jejuni Isolates Correlates with Lack of Shared Environments. Genome Biol Evol 2015;7(9):2663–2679 doi: 10.1093/gbe/evv174

35. Jolley KA, Bray JE, Maiden MCJ. Open-access bacterial population genomics: BIGSdb software, the PubMLST.org website and their applications. Wellcome Open Res 2018;3:124 doi: 10.12688/wellcomeopenres.14826.1

36. Mehat JW, La Ragione RM, van Vliet AHM. Campylobacter jejuni and Campylobacter coli autotransporter genes exhibit lineage-associated distribution and decay. BMC Genomics 2020;21(1):314 doi: 10.1186/s12864-020-6704-z

37. Dwivedi R, Nothaft H, Garber J, Xin Kin L, Stahl M et al. L-fucose influences chemotaxis and biofilm formation in Campylobacter jejuni. Mol Microbiol 2016;101(4):575–589 doi: 10.1111/mmi.13409

38. Seemann T. Prokka: rapid prokaryotic genome annotation. Bioinformatics 2014;30(14):2068–2069 doi: 10.1093/bioinformatics/btu153

39. Grissa I, Vergnaud G, Pourcel C. CRISPRFinder: a web tool to identify clustered regularly interspaced short palindromic repeats. Nucleic Acids Res 2007;35:W52–57 doi: 10.1093/nar/gkm360

40. Bland C, Ramsey TL, Sabree F, Lowe M, Brown K et al. CRISPR recognition tool (CRT): a tool for automatic detection of clustered regularly interspaced palindromic repeats. BMC Bioinformatics 2007;8:209 doi: 10.1186/1471-2105-8-209

41. Crooks GE, Hon G, Chandonia JM, Brenner SE. WebLogo: a sequence logo generator. Genome Res 2004;14(6):1188–1190 doi: 10.1101/gr.849004

42. Biswas A, Gagnon JN, Brouns SJ, Fineran PC, Brown CM. CRISPRTarget: Bioinformatic prediction and analysis of crRNA targets. RNA Biol 2013;10(5):817–827 doi: 10.4161/rna.24046

43. Gilchrist CLM, Chooi YH. Clinker & clustermap.js: Automatic generation of gene cluster comparison figures. Bioinformatics 2021:btab007 doi: 10.1093/bioinformatics/btab007

44. Rossi M, Silva M, Ribeiro-Goncalves BF, Silva DN, Machado MP et al. INNUENDO whole genome and core genome MLST schemas and datasets for Campylobacter jejuni (Version 1.0) [Data set]. Zenodo 2018:, doi: 10.5281/zenodo.1322564 doi:

45. Silva M, Machado MP, Silva DN, Rossi M, Moran-Gilad J et al. chewBBACA: A complete suite for gene-by-gene schema creation and strain identification. Microb Genom 2018;4(3):e000166 doi: 10.1099/mgen.0.000166

46. Zhou Z, Alikhan NF, Sergeant MJ, Luhmann N, Vaz C et al. GrapeTree: visualization of core genomic relationships among 100,000 bacterial pathogens. Genome Res 2018;28(9):1395–1404 doi: 10.1101/gr.232397.117

47. Kumar S. G. S, Tamura K. MEGA7: Molecular Evolutionary Genetics Analysis version 7.0 for bigger datasets. Mol Biol Evol 2016;33:1870–1874 doi: 10.1093/molbev/msw054

48. Marasini D, Karki AB, Bryant JM, Sheaff RJ, Fakhr MK. Molecular characterization of megaplasmids encoding the type VI secretion system in Campylobacter jejuni isolated from chicken livers and gizzards. Sci Rep 2020;10(1):12514 doi: 10.1038/s41598-020-69155-z

49. Dugar G, Leenay RT, Eisenbart SK, Bischler T, Aul BU et al. CRISPR RNA-Dependent Binding and Cleavage of Endogenous RNAs by the Campylobacter jejuni Cas9. Mol Cell 2018;69(5):893–905 doi: 10.1016/j.molcel.2018.01.032

50. Barrangou R. Nobel Dreams Come True for Doudna and Charpentier. CRISPR J 2020;3(5):317–318 doi: 10.1089/crispr.2020.29109.rba

51. Newire E, Aydin A, Juma S, Enne VI, Roberts AP. Identification of a Type IV-A CRISPR-Cas System Located Exclusively on IncHI1B/IncFIB Plasmids in Enterobacteriaceae. Front Microbiol 2020;11:1937 doi: 10.3389/fmicb.2020.01937

52. Kamruzzaman M, Iredell JR. CRISPR-Cas System in Antibiotic Resistance Plasmids in Klebsiella pneumoniae. Front Microbiol 2019;10:2934 doi: 10.3389/fmicb.2019.02934

53. Pinilla-Redondo R, Mayo-Munoz D, Russel J, Garrett RA, Randau L et al. Type IV CRISPR-Cas systems are highly diverse and involved in competition between plasmids. Nucleic Acids Res 2020;48(4):2000–2012 doi: 10.1093/nar/gkz1197

54. McDonald ND, Regmi A, Morreale DP, Borowski JD, Boyd EF. CRISPR-Cas systems are present predominantly on mobile genetic elements in Vibrio species. BMC Genomics 2019;20(1):105 doi: 10.1186/s12864-019-5439-1

55. Labbate M, Orata FD, Petty NK, Jayatilleke ND, King WL et al. A genomic island in Vibrio cholerae with VPI-1 site-specific recombination characteristics contains CRISPR-Cas and type VI secretion modules. Sci Rep 2016;6:36891 doi: 10.1038/srep36891

56. Xiao Y, Ng S, Nam KH, Ke A. How type II CRISPR-Cas establish immunity through Cas1-Cas2-mediated spacer integration. Nature 2017;550(7674):137–141 doi: 10.1038/nature24020

57. Pausch P, Al-Shayeb B, Bisom-Rapp E, Tsuchida CA, Li Z et al. CRISPR-CasPhi from huge phages is a hypercompact genome editor. Science 2020;369(6501):333–337 doi: 10.1126/science.abb1400

58. Pausch P, Soczek KM, Herbst DA, Tsuchida CA, Al-Shayeb B et al. DNA interference states of the hypercompact CRISPR-CasPhi effector. Nat Struct Mol Biol 2021;28(8):652–661 doi: 10.1038/s41594-021-00632-3

59. Hooton S, D’Angelantonio D, Hu Y, Connerton PL, Aprea G et al. Campylobacter bacteriophage DA10: an excised temperate bacteriophage targeted by CRISPR-cas. BMC Genomics 2020;21(1):400 doi: 10.1186/s12864-020-06808-3

60. Hooton SP, Connerton IF. Campylobacter jejuni acquire new host-derived CRISPR spacers when in association with bacteriophages harboring a CRISPR-like Cas4 protein. Front Microbiol 2014;5:744 doi: 10.3389/fmicb.2014.00744

61. Kovanen SM, Kivisto RI, Rossi M, Hanninen ML. A combination of MLST and CRISPR typing reveals dominant Campylobacter jejuni types in organically farmed laying hens. J Appl Microbiol 2014;117(1):249–257 doi: 10.1111/jam.12503

62. Louwen R, Horst-Kreft D, de Boer AG, van der Graaf L, de Knegt G et al. A novel link between Campylobacter jejuni bacteriophage defence, virulence and Guillain-Barre syndrome. Eur J Clin Microbiol Infect Dis, Research Support, Non-U.S. Gov’t 2013;32(2):207–226 doi: 10.1007/s10096-012-1733-4

63. Saha C, Horst-Kreft D, Kross I, van der Spek PJ, Louwen R et al. Campylobacter jejuni Cas9 Modulates the Transcriptome in Caco-2 Intestinal Epithelial Cells. Genes (Basel) 2020;11(10):1193 doi: 10.3390/genes11101193

64. Saha C, Mohanraju P, Stubbs A, Dugar G, Hoogstrate Y et al. Guide-free Cas9 from pathogenic Campylobacter jejuni bacteria causes severe damage to DNA. Sci Adv 2020;6(25):eaaz4849 doi: 10.1126/sciadv.aaz4849

65. Shabbir MA, Wu Q, Shabbir MZ, Sajid A, Ahmed S et al. The CRISPR-cas system promotes antimicrobial resistance in Campylobacter jejuni. Future Microbiol 2018;13:1757–1774 doi: 10.2217/fmb-2018-0234

66. Shabbir MAB, Tang Y, Xu Z, Lin M, Cheng G et al. The Involvement of the Cas9 Gene in Virulence of Campylobacter jejuni. Front Cell Infect Microbiol 2018;8:285 doi: 10.3389/fcimb.2018.00285

67. Corcionivoschi N, Gundogdu O, Moran L, Kelly C, Scates P et al. Virulence characteristics of hcp (+) Campylobacter jejuni and Campylobacter coli isolates from retail chicken. Gut Pathog 2015;7:20 doi: 10.1186/s13099-015-0067-z

68. Liaw J, Hong G, Davies C, Elmi A, Sima F et al. The Campylobacter jejuni Type VI Secretion System Enhances the Oxidative Stress Response and Host Colonization. Front Microbiol 2019;10:2864 doi: 10.3389/fmicb.2019.02864

69. Brown HL, Reuter M, Hanman K, Betts RP, van Vliet AHM. Prevention of biofilm formation and removal of existing biofilms by extracellular DNases of Campylobacter jejuni. PLoS One 2015;10(3):e0121680 doi: 10.1371/journal.pone.0121680

