## Supplementary Figures S1, S2, S3, S4 for "A *Campylobacter* integrative and conjugative element with a CRISPR-Cas9 system targeting competing plasmids: a history of plasmid warfare?"

#### LEGENDS TO SUPPLEMENTARY INFORMATION

**Figure S1.** Alignment of the native *C. jejuni* and *C. coli* Cas9 proteins with the CampyICE1 Cas9 proteins from *C. jejuni* and *C. coli* clades. Part A shows an alignment of the *C. jejuni* chromosomal Cas9 protein with the CampyICE1 Cas9 from a *C. jejuni* strain, whereas Part B combines the different version from the *C. jejuni* and *C. coli* clades. Asterisks indicate identical residues, full stops and colon indicate conservative substitutions. Functional domains of Cas9 are indicated in yellow shading, while conserved amino acid residues are highlighted by underlined bold typeface [34].

**Figure S2.** Genetic structure of CampyICE1 and related mobile genetic elements (integrative conjugative elements) present in related *Campylobacter* species, presented as output of a comparison of Prokka-generated annotations [38] using Clinker [43].

**Figure S3.** Graphical representation of genetic variability of the mobile elements CJIE1, CJIE2, CJIE3 and CJIE4 from *C. jejuni* and *C. coli* genomes, presented as output of a comparison of Prokka-generated annotations [38] using Clinker [43].

**Figure S4.** Graphical representation of genetic variability of the plasmids pCC42, pTet and pVir from *C. jejuni* and *C. coli* genomes, presented as output of a comparison of Prokka-generated annotations [38] using Clinker [43].

**van Vliet et al, Figure S1**

Part A

```
                       RuvC I                               R-rich region
Cj_Cas9       VARILAFDIGISSIGWAFSENDELKDCGVRIFTKVENPKTGESLALPRRLARSARKRLAR
Cj_ICE1_Cas9  -MKIIGFNLGIANIGWALRENDEIIDCGVRVFDIPENPKNGNSLALERRENKARMKIVKR
                :*:.*::**:.****: ****: *****:*   ****.*:**** **  ::  * : *

Cj_Cas9       RKARLNHLKHLIANEFKLNYEDYQSFDESLAKAYKGSLISPYELRFRALNELLSKQDFAR
Cj_ICE1_Cas9  KKARMLATKTFLKKELNVDLS---------KLFLIGSTQSIYELRTKALSSLISKEELSA
              :***:   * :: :*:::: .              **   * **** :**..*:**::::

Cj_Cas9       VILHIAKRRGYDDIKNSDDKEKGAILKAIKQNEEKLANYQSVGEYLYKEYFQKFKENSKE
Cj_ICE1_Cas9  IILHIAKHRGYDDSALKN--ENGTIIEALNKNKEAMLKFKSVGEYFYKNFVQ-----NKE
              :******:*****   .:  *:*:*::*:::*:* : :::*****:**::.*     .**

Cj_Cas9       FTNVRNKKESYERCIAQSFLKDELKLIFKKQREFGFSFSKKFEEEVLSVAFYKRALKDFS
Cj_ICE1_Cas9  VKKIRNTTEDYSNSVPRSLLKQELDLILDKQKELGLIKNADFKAKLFEIIFFKRPLKDFS
              ..::**..*.*...:.:*:**:**.**:.**:*:*:  . .*: :::.: *:**.*****

Cj_Cas9       HLVGNCSFFTDEKRAPKNSPLAFMFVALTRIINLLNNLKNTEGILYTKDDLNALLNEVLK
Cj_ICE1_Cas9  NKIGNCIFFENEKRAAKNTISACEFVALGKVVNLLKSIEKDIGIVYEKDSINEIMSIILD
              : :*** ** :****.**:  *  **** :::***:.:::  **:* **.:* ::. :*.

Cj_Cas9       NGTLTYKQTKKLLGLSDDYEFK-------GEKGTYFIEFKKYKEFIKALGE--HNLSQDD
Cj_ICE1_Cas9  KTSISYKKIRDILNLPQDINFKGLDYSKNNAENSKLVDLKKLNEFKKALGDGFTNLDKDI
              : :::**: :.:*.*.:* :**       . :.: ::::** :** ****:   **.:*

Cj_Cas9       LNEIAKDITLIKDEIKLKKALAKY-DLNQNQIDSLSKLEFKDHLNISFKALKLVTPLMLE
Cj_ICE1_Cas9  LDSIATDITLTKDTATLKEKLKNYNVLNAEQIEKLSELVFNDHINLSLKALKQIIPLMYE
              *:.**.**** **  .**: * :*  ** :**:.**:* *:**:*:*:**** : *** *

Cj_Cas9       GKKYDEACNELNLKVAINEDKKDFLPAFNETYYKDEVTNPVVLRAIKEYRKVLNALLKKY
Cj_ICE1_Cas9  GKRYDEACELCNFTIAKNQEKSEYLPLFEKTRFAKDISSPVVIRAICEFRKLLNDIIRRY
              **:*****:  *:.:* *::*.::** *::* : .:::.***:*** *:**:** ::::*

                   RuvC II
Cj_Cas9       GKVHKINIELAREVGKNHSQRAKIEKEQNENYKAKKDAELECEKLGLKINSKNILKLRLF
Cj_ICE1_Cas9  GSVHKIHLELTRDFGISFSDRKKIIKEIEQNEQSRIKALETIKELKLEETSKNIQIVRLF
              *.****::**:*:.* ..*:* ** ** ::* ::: .*    ::* *: .****  :***

                                                HNH domain
Cj_Cas9       KEQKEFCAYSGEKIKISDLQDEKMLEIDHIYPYSRSFDDSYMNKVLVFTKQNQEKLNQTP
Cj_ICE1_Cas9  EDQKGICPYSGLKMDLNRLDE---LVIDYIRPYNRSLDDSYSNKILTFKKLSDLKQGKTP
              ::** :*.*** *:.:. *::   * **:* **.**:**** **:*.*.* .: * .:**

Cj_Cas9       FEAFGNDSAKWQKIEVLAKNLPTKKQKRILDKNYKDKEQKNFKDRNLNDTRYIARLVLNY
Cj_ICE1_Cas9  FEAFGEDEKLWAEINERIKEYNGKKRFKIFDKFFKDKKPFDFTEQTLQDTRWLTKLVASY
              *****:*.  * :*:   *:   **: :*:** :***:  :*.::.*:***::::** .*

                                                                  RuvC III
Cj_Cas9       TKDYLDFLPLSDDENTKLNDTQKGSKVHVEAKSGMLTSALRHTWGFSAKDRNNHLHHAID
Cj_ICE1_Cas9  LNEYLSFLPISEDENTALGYGEKGSKQHVVLSSGMITQMLRNFWYLGFKNHKDYKNNAMD
               ::**.***:*:**** *.  :**** **  .***:*. **: * :. *::::: ::*:*

Cj_Cas9       AVIIAYANNSIVKAFSDFKKEQESNSAELYAKKISELDYKNKRKFFEPFSGFRQKVLDKI
Cj_ICE1_Cas9  AIIVAFTTNSIIFAFNNFKKELDLAKAEFYANKISESDYLLKRKFLPPFSGFKEQALEKV
              *:*:*::.***: **.:**** :  .**:**:**** **  ****: *****:::.*:*:

Cj_Cas9       DEIFVSKPERKKPSGALHEETFRKEEEFYQSYGGKEGVLKALELGKIRKVNGKIVK--NG
Cj_ICE1_Cas9  KNIFVSHSLKIKNKGTLHELTPLKIKELKNTYGDLD---LAVKLGKIRKYNDKYYANANG
              .:****:. : * .*:*** *  * :*: ::**. :    *::****** *.*     **
```

```
Cj_Cas9       DMFRVDIFKHKKTNKFYAVPIYTMDFALKVLPNKAVARSKKGEIKDWILMDENYEFCFSL
Cj_ICE1_Cas9  SLVRADLFVDKK-NKFHAVSIYKADFSTKKLPNKTPATTSNGETKEGIEMNENYNFCMSL
              .:.*.*:* .** ***:**.**. **: * ****: * :.:** *: * *:***:**:**

Cj_Cas9       YKDSLILIQTKDMQEPEFVYYNAFTSSTVSLIVSKHDNKFETLSKNQKILFKNANEKEVI
Cj_ICE1_Cas9  YKNTPISVKIKGMKEPIICYYHGFNTSGSKITYKKHDNNYHNLSEDEMVVFR-KNDK---
              **:: * :: *.*:** : **:.*.:*  .:  .****::..**::: ::*:  *:*

Cj_Cas9       AKSIGIQNLKVFEKYIVSALGEVTKAEFRQREDFKK
Cj_ICE1_Cas9  -ESIAVGKILEIKKYSISPSGELSLIENEERKWF--
               :**.: ::  ::** :*. **::  * .:*: *
```

Part B

```
                                      RuvC I
C_jejuni_Cas9                 VARILAFDIGISSIGWAFSENDELKDCGVRIFTKVENPKTGESLALPRRL
C_coli_Clade1a_Cas9           MARILAFDIGISSIGWAFSENDELKDCGVRIFTKAENPKTGESLALPRRL
C_coli_Clade1b_Cas9           MARILAFDIGISSIGWAFSENDELKDCGVRIFTKAENPKTGESLALPRRL
C_coli_Clade1c_Cas9           -MKILGFDIGINSIGWAFVEDNQLQDCGVRLFTKAEDPKTKESLALPRRN
C_coli_Clade2_Cas9            -MKILGFDIGINSIGWAFVEDNQLQDCGVRLFTKAEDPKRKESLALPRRN
C_coli_Clade3_Cas9            -MKILGFDIGINSIGWAFVEDNQLQDCGVRLFTKAEDPKTRESLALPRRN
C_jejuni_ICE1_Cas9            -MKIIGFNLGIANIGWALRENDEIIDCGVRVFDIPENPKNGNSLALERRE
C_coli_ICE1_Clade1a_Cas9      -MKIIGFNLGIANIGWALRENDEIIDCGVRVFDIPENPKNGNSLALERRE
C_coli_ICE1_Clade1c_Cas9      -MKILGFNLGIANIGWALRVNDEIIDCGVRVFDIPENPKNGNSLALERRE
C_coli_ICE1_Clade2_Cas9       -MKILGFNLGIANIGWALRENDEIIDCGVRVFDIPENPKNGNSLALERRE
C_coli_ICE1_Clade3_Cas9       -MKILGFNLGIANIGWALRVNDEIIDCGVRVFDIPENPKNGNSLALERRE
                                :*:.*::** .****:  :::: *****:*   *:**  :**** **

                               R-rich region
C_jejuni_Cas9                 ARSARKRLARRKARLNHLKHLIANEFKLNYEDYQSFDESLAKAYKGSLIS
C_coli_Clade1a_Cas9           ARSARKRLARRKARLNHLKHLIANEFKLNYEDYQSFDESLAKAYKGSLIS
C_coli_Clade1b_Cas9           ARSARKRLARRKARLNHLKHLIANEFKLNYEDYQSFDESLAKAYKGSLIS
C_coli_Clade1c_Cas9           ARSNRRRLGRRRSRLIALKHIISKELKLNYQDYMANDRELPKAYEGRLIS
C_coli_Clade2_Cas9            ARSNRRRLGRRRSRLIALKHIISKELKLNYQDYIANDGELPKAYEGRLVS
C_coli_Clade3_Cas9            ARSNRRRLGRRRSRLIALKHIISKELKLNYQDYIANDGELPKAYEGRLVS
C_jejuni_ICE1_Cas9            NKARMKIVKRKKARMLATKTFLKKELNVDLSKLFLIGS---------TQS
C_coli_ICE1_Clade1a_Cas9      NKARMKIVKRKKARMLATKTFLKKELNVDLSKLFLIGS---------TQS
C_coli_ICE1_Clade1c_Cas9      NKARMKTIKRKKARMLTTKTFLKKEFGIDLSKLFLIGS---------TQS
C_coli_ICE1_Clade2_Cas9       NKARMKTIKRKKARMLTTKTFLKKEFGIDLSKLFLIGS---------TQS
C_coli_ICE1_Clade3_Cas9       NKARMKTIKRKKARMLTTKTFLKKEFGIDLSKLFLIGS---------TQS
                               ::  : : *:::*:   * :: :*: :: ..    .            *

C_jejuni_Cas9                 PYELRFRALNELLSKQDFARVILHIAKRRGYDD--IKNSDDKEKGAILKA
C_coli_Clade1a_Cas9           PYELRFRALNELLSKQDFARVILHIAKRRGYDD--IKNNGDEEKSEILKA
C_coli_Clade1b_Cas9           PYELRFRALNELLSKQDFARVILHIAKRRGYDD--IKNSDDKEKGAILKA
C_coli_Clade1c_Cas9           PYELRYKALNEKIEPKDLARVILHIAKHRGYMNKNEKKSNDNEKGKILSA
C_coli_Clade2_Cas9            PYELRYKALNEKIEPKDLARVILHIAKHRGYMNKNEKKSSDNEKGKILSA
C_coli_Clade3_Cas9            PYELRYKALNEKIEPKDLARVILHIAKHRGYMNKNEKKSSDNEKGKILSA
C_jejuni_ICE1_Cas9            IYELRTKALSSLISKEELSAIILHIAKHRGYDDSALKN----ENGTIIEA
C_coli_ICE1_Clade1a_Cas9      IYELRTKALSSLISKEELSAIILHIAKHRGYDDSALKN----ENGTIIEA
C_coli_ICE1_Clade1c_Cas9      IYELRTKALNSLISKEELAAIILHIAKHRGYDDSALKN----ENGIIIEA
C_coli_ICE1_Clade2_Cas9       IYELRTKALNSLISKEELAAIILHIAKHRGYDDSALKN----ENGIIIEA
C_coli_ICE1_Clade3_Cas9       IYELRTKALNSLISKEELAAIILHIAKHRGYDDSALKN----ENGIIIEA
                               **** :**.. :. :::: :******:*** :   *:    *:. *:.*

C_jejuni_Cas9                 IKQNEEKLANYQSVGEYLYKEYFQKFKENSKEFTNVRNKKESYERCIAQS
C_coli_Clade1a_Cas9           IKQNEEKLVNYQSVGEYLYKEYFQKFKENSKEFINVRNKKESYERCIAQS
C_coli_Clade1b_Cas9           IKQNEEKLANYQSVGEYLYKEYFQKFKENSKEFTNVRNKKESYERCIAQS
C_coli_Clade1c_Cas9           LKTNALKLEKYQSVGEYFYKEFFQKHKENTKDFINIRNKKGSYDNCVLAS
C_coli_Clade2_Cas9            LKTNALKLEKYQSVGEYFYKEFFQKYKENTKDFINIRNKEGSYENCVLAS
C_coli_Clade3_Cas9            LKTNALKLEKYQSVGEYFYKEFFQKYRENTKDFINIRNKEGSYENCVLAS
C_jejuni_ICE1_Cas9            LNKNKEAMLKFKSVGEYFYKNFVQN-----KEVKKIRNTTEDYSNSVPRS
C_coli_ICE1_Clade1a_Cas9      LNKNKEAMLKFKSVGEYFYKNFVQN-----KEVKKIRNTTEDYSNSVPRS
C_coli_ICE1_Clade1c_Cas9      LNKNKEEMLKFKSVGEYFYKNFVQS-----KEVRKIRNTTEDYSNSIPRS
C_coli_ICE1_Clade2_Cas9       LNKNKEEMLKFKSVGEYFYKNFVQS-----KEVRKIRNTTEDYSNSIPRS
C_coli_ICE1_Clade3_Cas9       LNKNKEEMLKFKSVGEYFYKNFVQS-----KEVRKIRNTTEDYSNSIPRS
                              :: *   : :::*****:**::.*.     *:. ::**.  .*...:  *
```

```
C_jejuni_Cas9                 FLKDELKLIFKKQREFGFSFSKKFEEEVLSVAFYKRALKDFSHLVGNCSF
C_coli_Clade1a_Cas9           FLKDELKLIFQKQREFGFSFSKKFEEEVLSVAFYKRALKDFSHLVGNCSF
C_coli_Clade1b_Cas9           FLKDELKLIFKKQREFGFSFSKKFEEEVLSVAFYKRALKDFSHLVGNCSF
C_coli_Clade1c_Cas9           DLEKELKLILEKQKEWGYSYDDNFIKEILKVAFFQRPLKDFSYLVGVCTF
C_coli_Clade2_Cas9            DLEKELKLILEKQKEWGYSYNDNFIKEILKVAFFQRPLKDFSYLVGACTF
C_coli_Clade3_Cas9            DLEKELKLILEKQKEWGYSYNDNFIKEILKVAFFQRPLKDFSYLVGACTF
C_jejuni_ICE1_Cas9            LLKQELDLILDKQKELGLIKNADFKAKLFEIIFFKRPLKDFSNKIGNCIF
C_coli_ICE1_Clade1a_Cas9      LLKQELDLILDKQKELGLIKNADFKEKLFEIIFFKRPLKDFSNKIGNCIF
C_coli_ICE1_Clade1c_Cas9      LLKQELDLILNKQKELGLIKNADFKEKLFEIIFFKRPLKDFSNKIGNCIF
C_coli_ICE1_Clade2_Cas9       LLKQELDLILNKQKELGLIKNADFKEKLFEIIFFKRPLKDFSNKIGNCIF
C_coli_ICE1_Clade3_Cas9       LLKQELDLILNKQKELGLIKNADFKEKLFEIIFFKRPLKDFSNKIGNCIF
                               *:.**.**:.**:* *   . .*  :::.: *::*.*****  :* * *

C_jejuni_Cas9                 FTDEKRAPKNSPLAFMFVALTRIINLLNNLKNTEGILYTKDDLNALLNEV
C_coli_Clade1a_Cas9           FTDEKRAPKNSPLAFMFVALTRIINLLNNLKNTEGILYTKDDLNALLNEV
C_coli_Clade1b_Cas9           FTDEKRAPKNSPLAFMFVALTRIINLLNNLKNTEGILYTKDDLNALLNEV
C_coli_Clade1c_Cas9           FEDEKRACKNSYSAWEFVALTKIINELKSLEKESGELVSSQIINEILNHV
C_coli_Clade2_Cas9            FEDEKRACKNSYSAWEFVALTKIINELKSLEKESGELVSSQIINEILNHI
C_coli_Clade3_Cas9            FEDEKRACKNSYSAWEFVALTKIINELKSLEKESGELVSSQIINEILNHI
C_jejuni_ICE1_Cas9            FENEKRAAKNTISACEFVALGKVVNLLKSIEKDIGIVYEKDSINEIMSII
C_coli_ICE1_Clade1a_Cas9      FENEKRAAKNTISACEFVALGKVVNLLKSIEKDIGIVYEKDSINEIMSII
C_coli_ICE1_Clade1c_Cas9      FENEKRAAKNTLSACEFVALGKVINLLKSIEKDTGIVYEKDIISEIMNII
C_coli_ICE1_Clade2_Cas9       FENEKRAAKNTLSACEFVALGKVINLLKSIEKDTGIVYEKDIISEIMNII
C_coli_ICE1_Clade3_Cas9       FENEKRAAKNTLSACEFVALGKVINLLKSIEKDTGIVYEKDIISEIMNII
                              * :**** **:  *  **** :::* *:.:::  * :  .: :. ::. :

C_jejuni_Cas9                 LKNGTLTYKQTKKLLGLSDDYEFKG-------EKGTYFIEFKKYKEFIKA
C_coli_Clade1a_Cas9           LKNGTLTYKQTKKLLGLSDDYEFKG-------EKGTYFIEFKKYKEFIKA
C_coli_Clade1b_Cas9           LKNGTLTYKQTKKLLGLSDDYEFKG-------EKGTYFIEFKKYKEFIKA
C_coli_Clade1c_Cas9           LDKGSITYKKFREYIKLHESMKFKSLKYDKDNAESTKLIEFRKLVEFKKA
C_coli_Clade2_Cas9            LDKGSITYKKFREYIKLHESMKFKSLKYDKDNAESVKLIEFRKLVEFKKA
C_coli_Clade3_Cas9            LDKGSITYKKFREYIKLHESMKFKSLKYDKDNVESTKLIEFRKLVEFKKA
C_jejuni_ICE1_Cas9            LDKTSISYKKIRDILNLPQDINFKGLDYSKNNAENSKLVDLKKLNEFKKA
C_coli_ICE1_Clade1a_Cas9      LDKTSISYKKIRDILNLPQDINFKGLDYSKNNAENSKLVDLKKLNEFKKA
C_coli_ICE1_Clade1c_Cas9      LNKASISYKKIRDILNLPQDISFKGLDYSKNNAENSKLVDFKKLNEFKKA
C_coli_ICE1_Clade2_Cas9       LNKASISYKKIRDILNLPQDISFKGLDYSKNNAENSKLVDFKKLNEFKKA
C_coli_ICE1_Clade3_Cas9       LNKASISYKKIRDILNLPQDISFKGLDYSKNNAENSKLVDFKKLNEFKKA
                              *.: :::**: :. : * :. .**.        :.  :::::*  ** **

C_jejuni_Cas9                 LGEH--NLSQDDLNEIAKDITLIKDEIKLKKALAKY-DLNQNQIDSLSKL
C_coli_Clade1a_Cas9           LGEH--NLSQDDLNEIAKDITLIKDEIKLKKALAKY-DLNQNQIDSLSKL
C_coli_Clade1b_Cas9           LGDH--SLSQDDLNEIAKDITLIKDEIKLKKALAKY-DLNQNQIDSLSKL
C_coli_Clade1c_Cas9           LGEH--SLTREELDQIATYITLIKDNEKLKITLEKY-SLNNEQIKNLIEI
C_coli_Clade2_Cas9            LGEH--SLTREELDQIATYITLIKDNEKLKITLEKY-SLNNEQIKNLIEI
C_coli_Clade3_Cas9            LGEH--SLSREELDQIATYITLIKDNEKLKITLEKY-SLNNEQIKNLLEI
C_jejuni_ICE1_Cas9            LGDGFTNLDKDILDSIATDITLTKDTATLKEKLKNYNVLNAEQIEKLSEL
C_coli_ICE1_Clade1a_Cas9      LGDGFANLDKDILDSIATDITLTKDTATLKEKLKNYNVLNAEQIEKLSEL
C_coli_ICE1_Clade1c_Cas9      LDDSFVNLDKDILDSIATDITLTKDMTALKEKLESYNVLNKEQIEKLSEL
C_coli_ICE1_Clade2_Cas9       LDDSFVNLDKDILDSIATDITLTKDMTALKEKLESYNVLNKEQIEKLSEL
C_coli_ICE1_Clade3_Cas9       LDDSFVNLDKDILDSIATDITLTKDMTALKEKLESYNVLNKEQIEKLSEL
                              *.:   .* :: *:.**. *** **   **  * .*  ** :**..* ::

C_jejuni_Cas9                 EFKDHLNISFKALKLVTPLMLEGKKYDEACNELNLKVAINEDKKDFLPAF
C_coli_Clade1a_Cas9           EFKDHLNISFKALKLVTPLMLEGKKYDEACNELNLKVAINEDKKDFLPAF
C_coli_Clade1b_Cas9           EFKDHLNISFKALKLITPLMLEGKKYDEACNELNLKVAINEDKKDFLPAF
C_coli_Clade1c_Cas9           DFNDHINLSFKALNLILPLMKEGKRYDEGCKLLNLKTKSNNQKFDFLPAF
C_coli_Clade2_Cas9            DFNDHINLSFKALNLILPLMKEGKRYDEACKLLNLKTKSNNQKFDFLPAF
C_coli_Clade3_Cas9            DFNDHINLSFKALNLILPLMKEGKRYDEACKLLNLKTKSNNQKFDFLPAF
C_jejuni_ICE1_Cas9            VFNDHINLSLKALKQIIPLMYEGKRYDEACELCNFTIAKNQEKSEYLPLF
C_coli_ICE1_Clade1a_Cas9      VFNDHINLSLKALKQIIPLMYEGKRYDEACELCNFTIAKNQEKSEYLPLF
C_coli_ICE1_Clade1c_Cas9      AFNDYINLSLKALKQIIPLMYEGKRYDEACKLCNFAIAKNQEKSEFLPLF
C_coli_ICE1_Clade2_Cas9       AFNDYINLSLKALKQIIPLMYEGKRYDEACKLCNFAIAKNQEKSEFLPLF
C_coli_ICE1_Clade3_Cas9       AFNDYINLSLKALKQIIPLMYEGKRYDEACKLCNFAIAKNQEKSEFLPLF
                               *:*::*:*:***: : *** ***:***.*:  *:    *::* ::** *
```

```
                                                                     RuvC II
C_jejuni_Cas9                 NETYYKDEVTNPVVLRAIKEYRKVLNALLKKYGKVHKINIELAREVGKNH
C_coli_Clade1a_Cas9           NETYYKDEVTNPVVLRAIKEYRKVLNALLKKYGKVHKINIELAREVGKNH
C_coli_Clade1b_Cas9           NETYYKDEVTNPVVLRAIKEYRKILNALLKKYGKVHKINIELAREVGKNH
C_coli_Clade1c_Cas9           CDSIFAQELTNPIVNRAISEYRKVLNALLKKYGKMHKIHIELARDIGLSK
C_coli_Clade2_Cas9            CDSIFAQELTNPIVNRAISEYRKVLNALLKKYGKMHKIHIELARDIGLSK
C_coli_Clade3_Cas9            CDSIFAQELTNPIVNRAISEYRKVLNALLKKYGKMHKIHIELARDIGLSK
C_jejuni_ICE1_Cas9            EKTRFAKDISSPVVIRAICEFRKLLNDIIRRYGSVHKIHLELTRDFGISF
C_coli_ICE1_Clade1a_Cas9      EKTRFAKDISSPVVIRAICEFRKLLNDIIRRYGSVHKIHLELTRDFGISF
C_coli_ICE1_Clade1c_Cas9      EKTRFAKDISSPVVIRAVCEFRKLLNDIIRRYGSVHKIHLELTRDFGISF
C_coli_ICE1_Clade2_Cas9       EKTRFAKDISSPVVIRAVCEFRKLLNDIIRRYGSVHKIHLELTRDFGISF
C_coli_ICE1_Clade3_Cas9       EKTRFAKDISSPVVIRAVCEFRKLLNDIIRRYGSVHKIHLELTRDFGISF
                               .: : .:::.*:* **: *:**:** ::::**.:***::**:*:.* .

C_jejuni_Cas9                 SQRAKIEKEQNENYKAKKDAELECEKLGLKINSKNILKLRLFKEQKEFCA
C_coli_Clade1a_Cas9           SQRAKIEKEQNENYKAKKDAELECEKLGLKINSKNILKLRLFKEQKEFCA
C_coli_Clade1b_Cas9           SQRAKIEKEQNENYKAKKDAEIECEKLGLKINSKNILKLRLFKEQKEFCA
C_coli_Clade1c_Cas9           KLRTKIEKEQKENYENNIWALNECENFGLKANTKNILKLKLWKEQKEFCI
C_coli_Clade2_Cas9            KLRTKIEKEQKENYENNIWALNECENFGLKANAKNILKLKLWKEQKEFCI
C_coli_Clade3_Cas9            KLRAKIEKEQKENYENNIWALNECENFGLKANAKNILKLKLWKEQKEFCI
C_jejuni_ICE1_Cas9            SDRKKIIKEIEQNEQSRIKALETIKELKLEETSKNIQIVRLFEDQKGICP
C_coli_ICE1_Clade1a_Cas9      NDRKKIIKEIEQNEQSRIKALETIKELKLEETSKNIQIVRLFEDQKGICP
C_coli_ICE1_Clade1c_Cas9      NDRRKIIKEIEQNEQSRIKALETIKELKLEETPKNIQIVRLFGDQKGICP
C_coli_ICE1_Clade2_Cas9       NDRRKIIKEIEQNEQSRIKALETIKELKLEETPKNIQIVRLFGDQKGICP
C_coli_ICE1_Clade3_Cas9       NDRRKIIKEIEQNEQSRIKALETIKELKLEETPKNIQIVRLFGDQKGICP
                              . * ** ** ::* : .  *    ::: *: ..***  ::*: :** :*

                                                        HNH domain
C_jejuni_Cas9                 YSGEKIKISDLQDEKMLEIDHIYPYSRSFDDSYMNKVLVFTKQNQEKLNQ
C_coli_Clade1a_Cas9           YSGEKIKISDLQDEKMLEIDHIYPYSRSFDDSYMNKVLVFTKQNQEKLNQ
C_coli_Clade1b_Cas9           YSGEKIKLSDLQDEKMLEIDHIYPYSRSFDDSYMNKVLVFTKQNQEKLNK
C_coli_Clade1c_Cas9           YSGKKISIEHLRDEKTLEVDHIYPYSRSFDDSFLNKVLVFTKENQEKLNK
C_coli_Clade2_Cas9            YSGKKISIEHLRDEKALEVDHIYPYSRSFDDSFLNKVLVFTKENQEKLNK
C_coli_Clade3_Cas9            YSGKKISIEHLRDEKALEVDHIYPYSRSFDDSFLNKVLVFTKENQEKLNK
C_jejuni_ICE1_Cas9            YSGLKMDLNRLDE---LVIDYIRPYNRSLDDSYSNKILTFKKLSDLKQGK
C_coli_ICE1_Clade1a_Cas9      YSGLKMDLKCLDE---LVIDYIRPYNRSLDDSYSNKVLTFKKLNDLKQGK
C_coli_ICE1_Clade1c_Cas9      YSGLKMDLNRLDE---LVIDYIRPYNRSFDDSYSNKVLTFKKLNDLKQGK
C_coli_ICE1_Clade2_Cas9       YSGLKMDLNRLDE---LVIDYIRPYNRSFDDSYSNKVLTFKKLNDLKQGK
C_coli_ICE1_Clade3_Cas9       YSGLKMDLNRLDE---LVIDYIRPYNRSFDDSYSNKVLTFKKLNDLKQGK
                              *** *:.:. * :   * :*:* **.**:***: **:*.*.* .: * .:

C_jejuni_Cas9                 TPFEAFGNDSAKWQKIEVLAKNLPTKKQKRILDKNYKDKEQKNFKDRNLN
C_coli_Clade1a_Cas9           TPFEAFGNDSAKWQKIEVLAKNLPTKKQKRILDKNYKDKEQKNFKDRNLN
C_coli_Clade1b_Cas9           TPFEAFGNDSTKWQKIEVLAKNLPEKKQKRILDKNYKDKEQKDFKDRNLN
C_coli_Clade1c_Cas9           TPFEAFGANEERWSKIQALAQNLPYKKKNKILDEAFKGKQQQDFISRNLN
C_coli_Clade2_Cas9            TPFEAFGANEERWSKIQALAQNLPYKKKNKILDEAFKGKQQQDFISRNLN
C_coli_Clade3_Cas9            TPFEAFGANEERWSKIQALAQNLPYKKKNKILDEAFKGKQQQDFISRNLN
C_jejuni_ICE1_Cas9            TPFEAFGEDEKLWAEINERIKEYNGKKRFKIFDKFFKDKKPFDFTEQTLQ
C_coli_ICE1_Clade1a_Cas9      TPFEAFGEDEKLWAEINERIKEYNGKKRFKIFDKFFKDKKPFDFTEQTLQ
C_coli_ICE1_Clade1c_Cas9      TPFEAFGEDEKLWAEINERIKEYNGKKRFKIFDKFFKDKKPFDFTEQTLQ
C_coli_ICE1_Clade2_Cas9       TPFEAFGEDEKLWAEINERIKEYNGKKRFKIFDKFFKDKKPFDFTEQTLQ
C_coli_ICE1_Clade3_Cas9       TPFEAFGEDEKLWAEINERIKEYNGKKRFKIFDKFFKDKKPFDFTEQTLQ
                              ******* :.  * :*:   ::   **: :*:*: :*.*:  :* .:.*:

C_jejuni_Cas9                 DTRYIARLVLNYTKDYLDFLPLSDDENTKLNDTQKGSKVHVEAKSGMLTS
C_coli_Clade1a_Cas9           DTRYIARLVLNYTKDYLDFLPLSDDENTKLNDTQKGSKVHVEAKSGMLTS
C_coli_Clade1b_Cas9           DTRYIARLVLNYTKDYLDFLPLSDDENTKLNDTQKGSKVHVEAKSGMLTS
C_coli_Clade1c_Cas9           DTRYISTLIVKYTKEYLDFLPLDEKEDISLKSGEKGSKIHVQTINGMLTS
C_coli_Clade2_Cas9            DTRYISTLIVKYTKEYLDFLPLDEKEDINLKSGEKGSKIHVQTINGMLTS
C_coli_Clade3_Cas9            DTRYISTLIVKYTKEYLDFLPLDEKEDINLKSGEKGSKIHVQTINGMLTS
C_jejuni_ICE1_Cas9            DTRWLTKLVASYLNEYLSFLPISEDENTALGYGEKGSKQHVVLSSGMITQ
C_coli_ICE1_Clade1a_Cas9      DTRWLTKLVASYLNEYLSFLPISEDENTALGYGEKGSKQHVVLSSGMITQ
C_coli_ICE1_Clade1c_Cas9      DTRWLTKLVASYLNEYLLFLPISEDENTALGYGEKGSKQHVVLSSGMITQ
C_coli_ICE1_Clade2_Cas9       DTRWLTKLVASYLNEYLLFLPISKDENTALGYGEKGSKQHVVLSSGMITQ
C_coli_ICE1_Clade3_Cas9       DTRWLTKLVASYLNEYLLFLPISEDENTALGYGEKGSKQHVVLSSGMITQ
                              ***::: *: .* ::** ***:...*:  *   :**** **   .**:*.
                                               RuvC III
C_jejuni_Cas9                 ALRHTWGFSAKDRNNHLHHAIDAVIIAYANNSIVKAFSDFKKEQESNSAE
C_coli_Clade1a_Cas9           ALRHTWGFSAKDRNNHLHHAIDAVIIAYANNSIVKAFSDFKKEQESNSAE
C_coli_Clade1b_Cas9           ALRHTWGFSAKDRNNHLHHAIDAAIIAYANNSIVKAFSDFKKEQESNSVE
C_coli_Clade1c_Cas9           VLRHTWGFAQKDRNNHLHHALDATIVAYSTNAIIKAFSDFKKEQELLKAK
```

```
C_coli_Clade2_Cas9             VLRHTWGFAQKDRNNHLHHALDATIVAYSTNAIIKAFSDFKKEQELLKAK
C_coli_Clade3_Cas9             VLRHTWGFAQKDRNNHLHHALDATIVAYSTNAIIKAFSDFKKEQELLKAK
C_jejuni_ICE1_Cas9             MLRNFWYLGFKNHKDYKNNAMDAIIVAFTTNSIIFAFNNFKKELDLAKAE
C_coli_ICE1_Clade1a_Cas9       MLRNFWYLGFKNHKNYKNNAMDAIIVAFTTNSIIFAFNNFKKELDLAKAE
C_coli_ICE1_Clade1c_Cas9       MLRNFWYLGFKNYKDYKNNAMDAIIVAFTTNSIIFAFNNFKKELDLAKAE
C_coli_ICE1_Clade2_Cas9        MLRNFWYLGFKNYKDYKNNAMDAIIVAFTTNSIIFAFNNFKKELDLAKAE
C_coli_ICE1_Clade3_Cas9        MLRNFWYLGFKNYKDYKNNAMDAIIVAFTTNSIIFAFNNFKKELDLAKAE
                                **: * :. *: ::: ::*:** *:*::.*:*: **.:**** :  ..:

C_jejuni_Cas9                  LYAKKISELDYKNKRKFFEPFSGFRQKVLDKIDEIFVSKPERKKPSGALH
C_coli_Clade1a_Cas9            LYAKKISELDYKNKRKFFEPFSGFRQKVLDKIDEIFVSKPERKKPSGALH
C_coli_Clade1b_Cas9            LYAKKISELDYKNKRKFFEPFSGFRQKVLDKIDEIFVSKPERKKPSGALH
C_coli_Clade1c_Cas9            LYAKELTSDSYKHQAKFFEPFEGFREQILNQINKLFVSKPPRKRARGALH
C_coli_Clade2_Cas9             LYAKELTNDSYKHQAKFFEPFEGFREQILNQINKLFVSKPPRKRARGALH
C_coli_Clade3_Cas9             LYAKELTSDSYKHQAKFFEPFEGFREQILIQINKLFVSKPPRKRARGALH
C_jejuni_ICE1_Cas9             FYANKISESDYLLKRKFLPPFSGFKEQALEKVKNIFVSHSLKIKNKGTLH
C_coli_ICE1_Clade1a_Cas9       FYANKISESDYLLKRKFLPPFSGFKEQALEKVKNIFVSHSLKIKNKGTLH
C_coli_ICE1_Clade1c_Cas9       FYANKISESDYLLKRKFLPPFKGFKEQAFEKVNNIFVSHSLRIKNKGALH
C_coli_ICE1_Clade2_Cas9        FYANKISESDYLLKRKFLPPFKGFKEQAFEKVNNIFVSHSLRIKNKGALH
C_coli_ICE1_Clade3_Cas9        FYANKISESDYLLKRKFLPPFKGFKEQAFEKVNNIFVSHSLRIKNKGALH
                               :**::::. .*  : **: **.**::: : ::.::***:. : :  *:**

C_jejuni_Cas9                  EETFRKEEEFYQSYGGKEGVLKALELGKIRKVNGKIVKN--GDMFRVDIF
C_coli_Clade1a_Cas9            EETFRKEEEFYQSYGGKEGVLKALELGKIRKVNGKIVKN--GDMFRVDIF
C_coli_Clade1b_Cas9            EETFRKEEEFYQSYGGKEGVLKALELGKIRKVNGKIVKN--GDMFRVDIF
C_coli_Clade1c_Cas9            KETFYSKDEMIKKYNSQEGVEIALNCGKIRKIGTKYVEN--DTMVRIDIF
C_coli_Clade2_Cas9             KETFYSKDEMIKKYNSQEGVEIALNCGKIRKIGTKYVEN--DTMVRIDIF
C_coli_Clade3_Cas9             KETFYPKDEMIKKYNSQEGLEIALNCGKIRKIGTKYVEN--DTIVRIDIF
C_jejuni_ICE1_Cas9             ELTPLKIKELKNTYG---DLDLAVKLGKIRKYNDKYYANANGSLVRADLF
C_coli_ICE1_Clade1a_Cas9       ELTPLKIKELKNTYG---DLDLAVKLGKIRKYNDKYYVNAKGSLARTDLF
C_coli_ICE1_Clade1c_Cas9       NLNPLKIKELKNTYK---DLEFAIASGRIRNYNGKFYSNANGSLVRVDLF
C_coli_ICE1_Clade2_Cas9        NLNPLKIKELKNTYK---DLEFAIASGRIRNYNGKFYSNANGSLVRVDLF
C_coli_ICE1_Clade3_Cas9        NLNPLKIKELKNTYK---DLEFAIASGRIRNYNGKFYSNANGSLVRVDLF
                               : .    .*: :.*    .:  *:  *:**: . *   *  . : * *:*

C_jejuni_Cas9                  KHKKTNKFYAVPIYTMDFALKVLPNKAV--ARSKKGEIKDWILMDENYEF
C_coli_Clade1a_Cas9            KHKKTNKFYAVPIYTMDFALKVLPNKAV--ARSKKGEIKDWILMDENYEF
C_coli_Clade1b_Cas9            KHKKTNKFYAVPIYTMDFALKVLPNKAV--ARSKKGEIKDWILMDENYEF
C_coli_Clade1c_Cas9            KKQ--NKFYAIPIYTMDFALGVLPNKIVIAGKDKKGNPKQWQEIDESYEF
C_coli_Clade2_Cas9             KKQ--NKFYAIPIYTMDFALGVLPNKIVIIGKDKKGNPKQWQEIDESYEF
C_coli_Clade3_Cas9             KKQ--NKIYAIPIYTMDFALGVLPNKIVIKGKDKKGNPKQWQEIDESYEF
C_jejuni_ICE1_Cas9             VDK-KNKFHAVSIYKADFSTKKLPNKTP--ATTSNGETKEGIEMNENYNF
C_coli_ICE1_Clade1a_Cas9       VDK-ENKFHAVSIYKADFSTKKLPNKTP--ATTSNGETKEGIEMNENYNF
C_coli_ICE1_Clade1c_Cas9       VDK-KSKFHAVPIYKADFSTKQLPNKTP--AAISNGKIKEGVEMNENYNF
C_coli_ICE1_Clade2_Cas9        VDK-KSKFHAVPIYKADFSTKQLPNKTP--AAISNGKIKEGVEMNENYNF
C_coli_ICE1_Clade3_Cas9        VDK-KSKFHAVPIYKADFSTKQLPNKTP--AAISNGKIKEGVEMNENYNF
                                .:  .*::*:.**. **:   ****    .  .:*: *:   ::*.*:*

C_jejuni_Cas9                  CFSLYKDSLILIQTKDMQEPEFVYYNAFTSSTVSLIVSKHDNKFETLSKN
C_coli_Clade1a_Cas9            CFSLYKDSLILIQTKDMQEPEFVYYNAFTSSTVSLIVSKHDNKFETLSKN
C_coli_Clade1b_Cas9            CFSLYKDSLILIQTKDMQEPELVYFNAFTSSTVSLIVSKHDNKFETLSKN
C_coli_Clade1c_Cas9            CFSLHKDDLVLIQKKDMQEPEFAYYNGFDISNSSICVEKHDNKFENLTDN
C_coli_Clade2_Cas9             CFSLHKDDLVLIQKKDMQEPEFAYYNGFDISNSSICVEKHDNKFENLTDN
C_coli_Clade3_Cas9             CFSLHKDDLVLIQKKDMQEPEFAYYNSFDISNSSICVEKHDNKFENLTDN
C_jejuni_ICE1_Cas9             CMSLYKNTPISVKIKGMKEPIICYYHGFNTSGSKITYKKHDNNYHNLSED
C_coli_ICE1_Clade1a_Cas9       CMSLYKNTPVSVKIKGMKEPIICYYHGFNTSGSKITYKKHDNNYHNLSED
C_coli_ICE1_Clade1c_Cas9       YISLYKNTPISVKIKGMEEPIICYYHGFNISGSQIIYKKHDNNYYNLSKD
C_coli_ICE1_Clade2_Cas9        YISLYKNTPISVKIKGMEEPIICYYHGFNISGSQIIYKKHDNNYYNLSKD
C_coli_ICE1_Clade3_Cas9        YISLYKNTPISVKIKGMEEPIICYYHGFNISGSQIIYKKHDNNYYNLSKD
                                :**:*:  : :: *.*:** : *::.*  *  .:  .****:: .*:.:

C_jejuni_Cas9                  QKILFKNANEKEVIAKSIGIQNLKVFEKYIVSALGEVTKAEFRQREDFKK
C_coli_Clade1a_Cas9            QKILFKNANEKEVIAKSIGIQNLKVFEKYIVSALGEVTKAEFRQREDFKK
C_coli_Clade1b_Cas9            QKILFKNANEKEVIAKSIGIQNLKVFEKYIVSALGEVTKAEFRQREDFKK
C_coli_Clade1c_Cas9            QKLLFVNAKEGSVKAEKLGIQGLKIFEKYIITPLGEKIKADFKPREDIAL
C_coli_Clade2_Cas9             QKLLFTNAEEGNVKAKKIGIQGLRIFEKYIITPLGEKIKADFKPREDIAL
C_coli_Clade3_Cas9             QKLLFTNAEEGNVKAKKLGIQGLKIFEKYIITPLGEKIKADFKLREDIAL
C_jejuni_ICE1_Cas9             EMVVFRKNDKE-----SIAVGKILEIKKYSISPSGELSLIENEERKWF--
C_coli_ICE1_Clade1a_Cas9       EMVVFRKNDKE-----SIVVGKILEIKKYSISPSGELSLIENEKRKWF--
C_coli_ICE1_Clade1c_Cas9       EMAIFRKGDKE-----AIAIGRMLEIKKYNISPSGELILIENEERKWF--
C_coli_ICE1_Clade2_Cas9        EMAIFRKGDKE-----AIAIGRMLEIKKYNISPSGELILIENEERKWF--
```

```
C_coli_ICE1_Clade3_Cas9       EMAIFRKGDKE-----AIAIGRILEIKKYNISPSGELILIENEERKWF--
                              :  :* : .:       : :  :  ::** ::. **    : . *: :

C_jejuni_Cas9                 --------
C_coli_Clade1a_Cas9           --------
C_coli_Clade1b_Cas9           --------
C_coli_Clade1c_Cas9           KTSKKHGL
C_coli_Clade2_Cas9            KTSKKHGI
C_coli_Clade3_Cas9            KTSKKHGL
C_jejuni_ICE1_Cas9            --------
C_coli_ICE1_Clade1a_Cas9      --------
C_coli_ICE1_Clade1c_Cas9      --------
C_coli_ICE1_Clade2_Cas9       --------
C_coli_ICE1_Clade3_Cas9       --------
```

### ICE1-like elements in *Campylobacter*

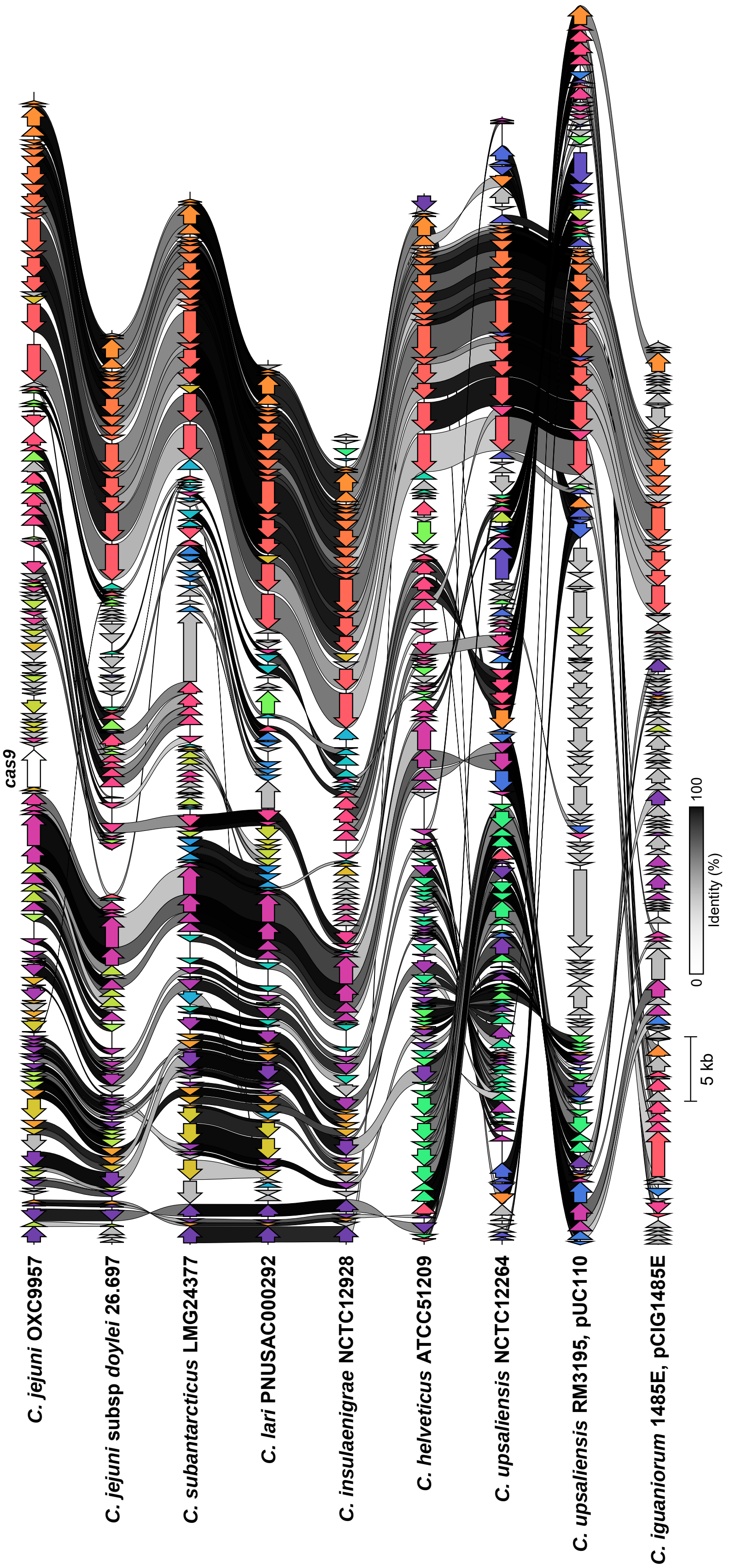


Van Vliet et al, Figure S2

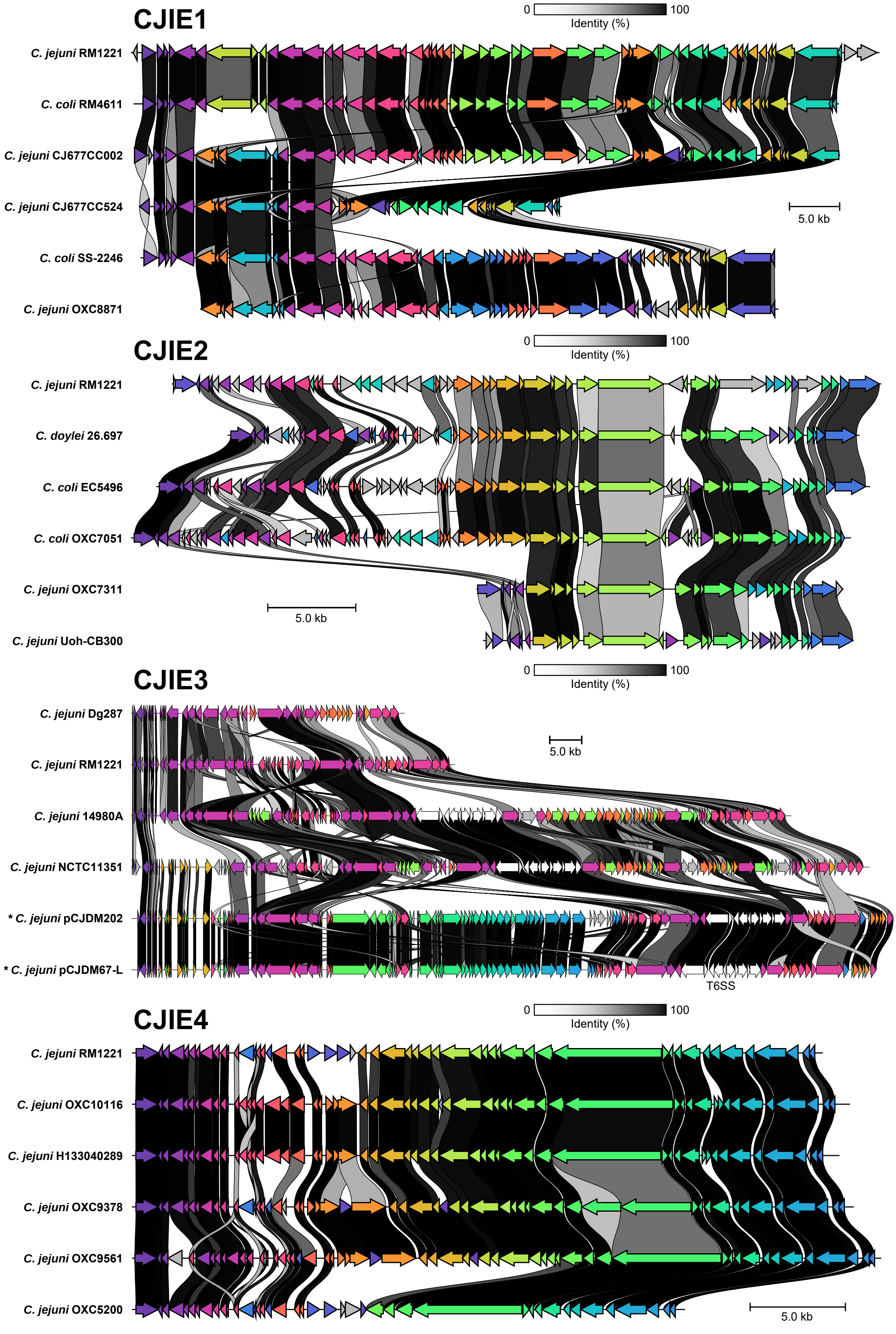


Van Vliet et al, Figure S3

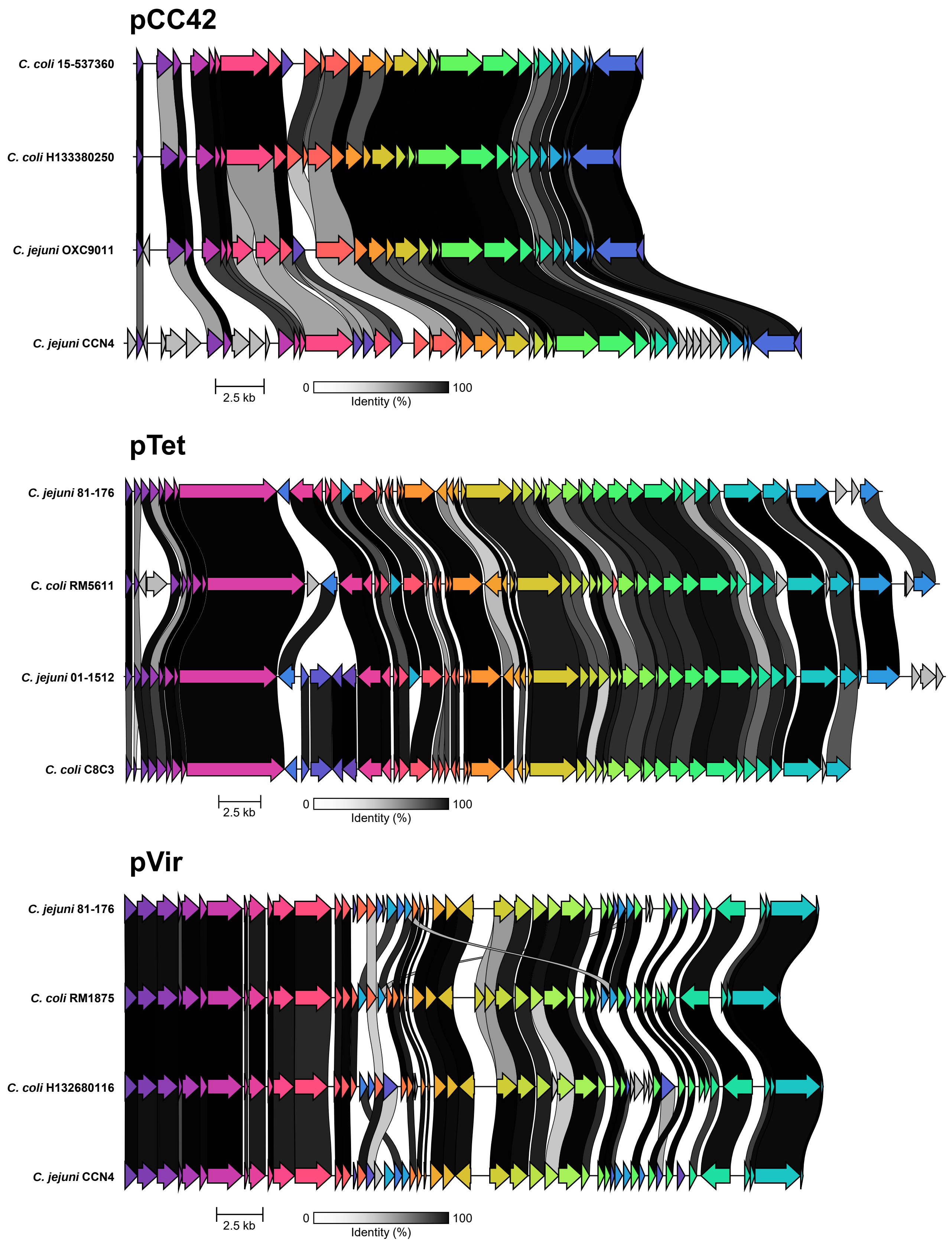


Van Vliet et al, Figure S4
