## Supplementary material for "A *Campylobacter* integrative and conjugative element with a CRISPR-Cas9 system targeting competing plasmids: a history of plasmid warfare?": Table S4

### LEGENDS TO SUPPLEMENTARY INFORMATION

**Table S1.** Overview of *C. jejuni* and *C. coli* genomes used in this study, with the Genbank or Campylobacter PubMLST accession numbers, metadata (source, MLST clonal complex, sequence type), and the presence/absence of the CJIE1-4 and CampyICE1 mobile elements and the pCC42, pTet and pVir plasmids. The position on the cgMLST tree used in Figures 3 and 4 is provided for ordering purposes [separate Excel spreadsheet].

**Table S2.** Sequences of CampyICE1 spacers and variant families with their predicted targets, as identified using CRISPRTarget [42] [separate Excel spreadsheet].

**Table S3.** Description of the three CampyICE1 spacer arrays per CampyICE1-positive *C. jejuni* or *C. coli* genome, and linkage to plasmid carriage [separate Excel spreadsheet].

**Table S4.** Distribution of CampyICE1 plasmid-specific CRISPR-spacers and pVir, pTet and pCC42 plasmids in CampyICE1-positive *C. jejuni* and *C. coli*.

**Table S4.** Distribution of CampyICE1 plasmid-specific CRISPR-spacers and pVir, pTet and pCC42 plasmids in CampyICE1-positive *C. jejuni* and *C. coli*.

| **plasmid** | **no plasmid** | | **plasmid present** | | |
|---|---|---|---|---|---|
| **spacer** [a] | **absent** | **matched** [b] | **absent** | **ΔCas9+mismatch**[b] | **matched** [b] |
| ***C. jejuni* (n=133)** | | | | | |
| pVir | 115 | 16 | 2 | 0+0 | 0 |
| pTet | 11 | 105 | 2 | 2+5 | 8 |
| pCC42 | 22 | 97 | 3 | 0+4 | 7 |
| ***C. coli* (n=81)** | | | | | |
| pVir | 27 | 50 | 3 | 1+0 | 0 |
| pTet | 8 | 58 | 3 | 1+11 | 0 |
| pCC42 | 2 | 57 | 0 | 2+9 | 11 |

a. Plasmid spacers identified by CRISPRfinder, CRISPR Recognition Tool CRT and manual searches were screened for matches with Campylobacter plasmids using CRISPRtarget.

b. CampyICE1-positive genomes positive for pVir, pTet and pCC42 were searched with the plasmid-targeting spacers using BLAST, and recorded for perfect matches and imperfect matches. This was to allow for possible sequence differences with the reference pVir, pTet and pCC42 plasmid sequences, or alternatively to detect mutations introduced to escape CRISPR-Cas functionality. In addition, the presence of a full-length *cas9* gene was checked, as this is required for CRISPR-Cas9 functionality.
